# Impaired Function in Diabetic Patient iPSCs-derived Blood Vessel Organoids Stem from a Subpopulation of Vascular Cells

**DOI:** 10.1101/2022.07.18.500478

**Authors:** Hojjat Naderi-Meshkin, Magdalini Eleftheriadou, Garrett Carney, Victoria A Cornelius, Clare-Ann Nelson, Sophia Kelaini, Andrew Yacoub, Clare Donaghy, Philip D Dunne, Raheleh Amirkhah, Anna Zampetaki, Lingfang Zeng, Alan W Stitt, David J Grieve, Andriana Margariti

## Abstract

The presence of both endothelial cells (ECs) and mural cells are central to the proper function of blood vessels in health and pathological changes in diseases including diabetes. Although iPSCs-derived vascular organoids (VOs) provide an appealing in vitro disease model and platform for drug screening, whether these organoids recapitulate human disease remains debatable. Here, we show human diabetic (DB)-VOs represent impaired vascular function including enhanced ROS activity, with higher mitochondrial content and activity, increased pro-inflammatory cytokines, and less regenerative potential in vivo. Using single-cell RNA sequencing, we identify all specialized types of vascular cells (artery, capillary, vein, lymphatic and tip cells, as well as pericytes and vSMCs) within vascular organoids, while demonstrating the dichotomy landscape of ECs and mural cells. Furthermore, we reveal basal heterogeneity within vascular organoids and demonstrate differences between diabetic and non-diabetic VOs. Of note, a subpopulation of ECs significantly enrich for ROS and oxidative phosphorylation hallmarks in DB-VOs, may represent early signs of aberrant angiogenesis in diabetes. This study helps to identify key biomarkers for diabetic disease progression and find signalling molecules amenable to drug intervention.

## 1. Introduction

Blood vessels deliver oxygen and nutrients to tissues and organs of the body. Improper function of blood vessels, referred to as cardiovascular diseases (CVDs), is the leading cause of death worldwide, representing 32% of all global deaths according to the World Health Organization ([WHO](https://www.who.int/news-room/fact-sheets/detail/cardiovascular-diseases-(cvds)#:~:text=Cardiovascular%20diseases%20(CVDs)%20are%20the,%2D%20and%20middle%2Dincome%20countries)). Two main cellular components of blood vessels are mural cells (pericytes and vascular smooth muscle cells (VSMCs)) and endothelial cells (ECs). Both mural cells and ECs are required for proper vascular function. Consequently, these cells are central to pathological dysfunctions including atherosclerosis, the most common CVDs, which can eventually damage not only the heart but also every single organ in the human body (Li *et al*., 2018).

Although many factors can lead to the development of CVDs, one stands out amongst the rest is Diabetes Mellitus (DM); a chronic metabolic disease that affects over 400 million people worldwide. DM increases two-fold the risk of developing CVDs as opposed to that of any individual not presenting this condition (Taylor *et al*., 2013). The traditional narrative linking both diseases centers around the presence of risk factors like hypertension, obesity, and dyslipidemia. However, the association is far more complex and may be due to alterations in both vascular cells’ composition and function. Recent studies have shown convincing evidence suggesting that high glucose concentration in the blood damages ECs and accelerates endothelial dysfunction (ED) (Domingueti *et al*., 2016; Giglio *et al*., 2021). Therefore, the implication is that DM potentiates ED, which subsequently becomes the driving force behind that leads to the genesis and progression of CVDs. The hyperglycemic state characteristic of uncontrolled DM promotes the development of CVDs; however, the relationship between both diseases is not yet fully understood.

Patient iPSCs-derived vascular organoids (VO) are the latest innovation for research and are being used as the quintessential model for *“in-vivo”* and *“in vitro”* studies. In particular, the presence of both mural cells and ECs in the vascular organoids are revealing new data that was previously unattainable. This information may prove vital in the fight against DM and CVDs. Previous studies have shown that iPSCs harvested from non-diabetic individuals can sprout normal blood vessels, however, the same does not hold for those from diabetic patients (Yang *et al*., 2020). Diabetic iPSCs seem to retain metabolic memory (Yu *et al*., 2012) or hyperglycemic memory (Luc *et al*., 2019) epigenetically, and as they differentiate into blood vessels, they become inflamed and are more prone to oxidative stress leading back to CVD (Su *et al*., 2021). Furthermore, studies have revealed the importance of paracrine signalling in controlling cellular heterogeneity and highlighted using multicellular populations such as organoids to better mimic native pathophysiological responses (Shalek *et al*., 2014). Additionally, vascular organoids exclude the complexity and variations that come from the organ itself or other non-vascular cells but have the essential vascular cells vital for blood vessel function allowing straightforward recapitulating and capturing pathophysiological changes *in vitro* (Kalucka *et al*., 2020; Gurevich *et al*., 2021; Pasut *et al*., 2021; Ricard *et al*., 2021). However, how disease-specific phenotypes are displayed in iPSCs-derived vascular organoids has not been fully delineated. Single-cell RNA sequencing provides an unprecedented opportunity to understand cell population changes in diabetic disease and find functional differences that are likely imprinted in the transcriptome and signalling pathways.

In this study, we have reproducibly made vascular organoids (VOs) from iPSCs reprogrammed from mononuclear cells (MNCs) of non-diabetic (NDs) and diabetic (DBs) individuals. Our immunofluorescence staining based on confocal microscopy and single-cell RNA-seq data showed the presence of vascular cells, mainly arteries and pericytes, in both ND-VOs and DB-VOs. We show convincing evidence confirming that diabetic organoids are more prone to produce pro-inflammatory cytokines and antiangiogenic proteins in conditioned media over time. Furthermore, diabetic organoids revealed significantly higher mod-LDL uptake and enhanced production of reactive oxygen specious (ROS), which was further confirmed by confocal images and single cell RNA sequencing. To the best of our knowledge, here for the first time, we analysed iPSCs-derived VOs of diabetic patients in comparison to non-diabetic individuals in a single-cell resolution. We identified 3 different subpopulations of ECs, one of which was enriched for hallmarks of MYC-TARGETs v1, oxidative phosphorylation (OXI-PHOS), and ROS production pathways in diabetic organoids, may represent early signs of aberrant angiogenesis in diabetes.

## 2. Material and methods

### 2.1. Experimental design

As illustrated in **Extended Data Fig. 1**, iPSCs were reprogrammed from MNCs of NDs and DBs individuals and differentiated into vascular cells in a 3D manner to get diabetic and non-diabetic vascular organoids, called DB-VOs and ND-VOs respectively. To better disease modelling of diabetic vasculopathy/angiopathy *in vitro*, established mature DB-VOs at day 15 from diabetic iPSCs were started to culture in high glucose concentration (33mM, adapted from our previous publication (Yang *et al*., 2020)) plus 1 ng/ml human TNF (Invitrogen, PHC3011) and 1 ng/ml IL-6 (Invitrogen, PHC3011)) (Wimmer, Leopoldi, Aichinger, Wick, *et al*., 2019). As a control, to maintain osmotic pressure the same in ND-VOs as DB-VOs, D-mannitol (Sigma-Aldrich) was added to the culture medium of ND-VOs to a final concentration of 33mM. D-mannitol is metabolically inert in humans and used as control of high glucose treatment experiments (Wimmer, Leopoldi, Aichinger, Wick, *et al*., 2019). DB-VOs and ND-VOs were exploited for deeper biological insights interrogated by both wet lab and dry lab analyses. For each analysis, at least three biological replicates were used and experiment-specific details have been mentioned accordingly. Statistical analyses were performed using Prism 9.3. 1 (GraphPad) or R 4.1.2. All statistical tests used are described in the Fig. legends. P < 0.05 or FDR < 0.05 was accepted as statistically significant.

### 2.2. Biological Materials and Ethical Considerations

Ethical approval of human peripheral blood samples from diabetic and nondiabetic donors to isolate MNCs and subsequently reprogramme them to iPSCs was from the Office for Research Ethics Committees of Northern Ireland (ORECNI) (REC 14/NI/1109). On the day of blood sample collection, a detailed medical history was obtained, and blood pressure was measured. A 20 ml sample of peripheral blood was obtained by venepuncture into VACUETTE® K3 EDTA-coated 4-ml tubes (454021, GREINER Bio-One). Verbal and written information about the study was provided to all participants and written informed consent was obtained before study procedures from those willing and consenting to take part. Patients with type 1 and type 2 diabetes of more than 15 years standing and age- and sex-matched non-diabetic volunteers, who acted as controls, were recruited for this study. Patients unable to provide informed consent for the study were excluded. All methods regarding the human iPSC culture and differentiation were performed following the relevant institutional guidelines and regulations including the Declaration of Helsinki and Ethical Guidelines for Medical and Health Research Involving Human Subjects.

Animals used in these studies were housed with free access to standard food and water at a room temperature of 21 ± 2 °C relative humidity of 45 ± 15% and a 12-h-light/dark cycle. All experiments were performed in accordance with the Guidance on the Operation of the Animals (Scientific Procedures) Act, 1986 and approved by the Queen’s University Belfast Animal Welfare and Ethical Review Body. Work was performed under the project license number PPL2922.

### 2.3. Reprogramming and characterization of human iPSCs

MNCs were reprogrammed from diabetic and non-diabetic donors (**Supplementary Table 1**) as previously described (Vilà-González *et al*., 2019). Briefly, 20 ml blood was collected into VACUETTE® K3 EDTA-coated 4 ml tubes (454021, GREINER Bio-one) and its MNCs fraction was separated by gradient centrifugation on Histopaque solution (10771, SIGMA) (1:1 ratio) for 30 min at 550g, RT with the brake off. The MNCs buffy coat was collected, after 3 washes with PBS, plated in an MNC medium, 4 million cells/ml. After expanding for 7 days, the cells were used for reprogramming to generate patient-specific iPSCs. Two million of MNCs were transfected with 10 μg of plasmid (5:1 ratio pEB-C5:pEBTg) as instructed by the Lonza CD34+ nucleofector kit (VPA-1003, Lonza) and Amaxa nucleofector (program T-016) according to the manufacturer’s protocol. Two days after transfection, the cells were cultured onto inactivated feeders (mouse embryonic fibroblasts from ATCC) in a reprogramming medium (KnockOut™ DMEM/F-12 with MEM Non-Essential Amino Acids Solution and GlutaMAX™ supplemented with KO Serum replacement (20%), 2-Mercaptoethanol (0.1 mM), all from Thermo Fisher Scientific, and human recombinant FGF2 (10 ng/ml), from Bio-Techne. Colonies of iPSCs appeared from day 9 within the reprogramming medium, which was subsequently sub-cultured once per week with a 1:6 ratio. The pluripotency capacity of the generated iPSC lines was verified before proceeding to the step of vascular organoid generation.

### 2.4. Vascular organoid generation

Blood vessel organoids were generated as previously described by Wimmer *et al*. 2019, with some critical modifications that dramatically decrease both labor-time and risk of contamination of vascular organoids. Briefly, iPSCs were seeded on a Cell-Matrix Coated 100mm cell culture dish and expanded for four days in StemMACS™ iPS Brew XF. When iPSCs reached 80-90% confluency, they were gently detached into cell clusters by 5 min incubation in StemMACS™ Passaging Solution XF. The cell cluster pellet was collected by centrifuge at 600 g, for 5 min. The pellet of the iPSC was distributed in small clamps (3-10 cells) into one full ultra-low attachment 6-well plate. The cell aggregates were generated within KO-DMEM plus 15 KO-Serum supplemented with ROCK inhibitor Y-27632 incubated for only 1 night. It is always critical to evenly distribute the cells/aggregates/spheroids inside Ultra-low attachment 6-well plates just before putting them in the 37ºC and 5% CO2 incubator.

After reaching a stable aggregate size (>50–200 μm in diameter), the aggregates were precipitated by gravitation for 20 min, and mesodermal induction was initiated in N2B27 neurobasal media containing CHIR99021 (12 µM) and BMP-4 (30 ng/mL) for 3 days. For another 2 days, spheroids were transferred into N2B27 neurobasal media containing Forskolin (2 µM) and hVEGF-A (100 ng/mL) to induce the budding of endothelial tubes. We then proceeded to collect the floating spheroids by gravitation and embedded them in a 3D collagen I–Matrigel matrix in 12-well plates (6500 µL per well) which allowed the sprouting of vascular networks. 3D collagen I–Matrigel matrix containing homogeneously distributed spheroids was left for 2 hours in 37 ºC and 5% CO_2_ incubator to let it polymerize, and then 1.5 mL of prewarmed StemPro34 (supplemented with VEGF-A (100 ng/mL) and hFGF2 (100 ng/mL) and 15% FBS) was gently poured onto each matrix from the 12-well wall and incubated for 3 days. From now on, the same media were exchanged every other day. After five days in the matrix, the sprouted vascular networks became visible enough to cut every 3-5 of them using a fine-tip, curved tweezer (**Supplementary Video 1**). The cut vascular networks were transferred back to the Ultra-low attachment 6-well plate gently by a 10 mL serological pipette, to let them self-assemble into mature organoids during the next 5 days. At this stage, they were ready to assess the presence of ECs and pericytes, or for any other experiment. For drug screening, every single mature organoid could be transferred into low-attachment U-bottom 96-well plates by a wide-bore blue tip.

### 2.5. Immunocytochemistry of vascular organoids

Vascular organoids (3-10) were transferred into 2 mL microtubes by a wide orifice blue tip (Alphalaboratories), washed once with PBS, and fixed in paraformaldehyde solution (4%) for 2 hours. Then they were incubated for 3 hours in blocking/permeabilization solution (3% FBS, 1% BSA (wt/vol), 0.5% Triton X-100, 0.5% Tween 20, and 0.01% (wt/vol) sodium deoxycholate in PBS (Wimmer, Leopoldi, Aichinger, Kerjaschki, *et al*., 2019)), 2 mL per microtube while shaking horizontally on a rocking shaker, after which they were incubated overnight at 4 °C with antibodies: ColIV (ab6311) for basement membrane, CD31 (Ab28364) for ECs and PDGFR-ß (AF385, R&D Systems) for pericytes. Samples were then washed 3 times with PBS and blocked in a 5% donkey serum in PBS solution, followed by washes, adding the appropriate secondary antibodies against the respected host of primary antibodies. incubating and counterstaining with DAPI (Life Technologies, D1306). The stained VOs were mounted on the iSpacers (Sunjin Lab) and mounting media was added at the end of the staining protocol. Samples were covered by a cover slip, examined under Confocal Microscopy (SP8, Leica), and their images were analyzed using ImageJ.

For live staining, VOs were incubated with Calcein-AM (Invitrogen, C3100) for 30 min, and immediately mounted on the iSpacers to capture images with Nikon 6D Live Cell Imaging Inverted Microscope within 2 hours.

### 2.6. Human protein array for both vascular organoids and their secreted media

After 21 days of treatment, protein extracts of VOs were collected and analysed using the human protein array (RayBiotech, AAH-ANG-1000-8). To use conditioned media for protein array investigation, at 42 days after treatment, 1.5 ml of culture media (minus FBS) was added onto each well of 6 well-plates containing 10 organoids, and conditioned media were collected after 24 hours. In total, 4.5 mL mixture of 3 DBs versus 4.5 mL of 3 NDs conditioned media were used for protein quantification, after a 2 min centrifugation at high speed to exclude any debris.

Quick Start™ Bradford protein assay was used to quantify and normalize protein concentration among samples. Additionally, internal positive and negative controls of each array membrane were used to normalize arrays together and subtract signal background, respectively, based on the RayBiotech instruction. Protein array membranes were exposed on chemiluminescence detection, and images were captured by G:BOX (Syngene). Protein Array plugin (Biii) in the ImageJ software was used to normalize and quantify protein levels with pixel intensity on the human protein array membranes containing 43 antibodies.

### 2.7. ROS production assay

Total Reactive Oxygen Species (ROS) Assay Kit 520 nm (Invitrogen, Cat: 88-5930) was used to identify ROS in the cells of vascular organoids by Flow cytometry in the FITC channel as well as Confocal imaging. Briefly, VOs were first broken into small pieces with 21-25G needle and then further digested into single cells in DLD for 20 min, cultured on 6-well plates for 3-4 days in the same media used for maintaining VOs in culture (StemPro34 medium with its supplements). After removing debris by washing with PBS, the single cells were incubated with the kit for 1hour as per the manufacturer’s instructions. Immediately, the cells were dissociated using TrypLE™ Express Enzyme (1X) (Thermo Fisher), filtered through pluriStrainer® 40 µm Mesh, and subjected to FACS florescent reading in PBS. Some small pieces were kept to image ROS fluorescent immediately by Nicon 6D Live imaging.

### 2.8. Uptake of acetylated-LDL

The capacity of lipid uptake was monitored using Alexa Fluor™594 AcLDL (Invitrogen, L35353). Similar to the ROS assay, dissociated VOs were cultured in a 6 well plate for 3-4 days, then incubated with 2.5 μg/ml Ac-LDL– Alexa Fluor™594 in StemPro34 medium for 4 hours at 37 °C immediately before Confocal or Flow Cytometry analysis. Thereafter, the cells were dissociated using TrypLE™ Express Enzyme (1X) (Thermo Fisher), filtered through pluriStrainer® 40 µm Mesh, and subjected to Flow Cytometry fluorescent reading in PBS. Small pieces of broken organoids were also used for Confocal imaging (SP8, Leica) of cellular uptake of Ac-LDL in parallel. Therefore, these VO pieces were further subjected to fixation with 4% PFA for 10 min and counterstained with DAPI.

### 2.9. Measuring mitochondrial content, number, and size

MitoTracker Red is a fluorescent dye taken up by active mitochondria based on their potential. This was used to quantify mitochondrial activity within VOs. VOs were placed in a low-attachment 96-well plate. MitoTracker Red CMXROS (Invitrogen, M7512) was made up to 500nM in organoid culture media. Organoids were incubated for 1 hour and 30 minutes at 37°C and 5% CO2. The fluorescence was measured using the OMEGA plate reader (average of 20 reads per well). Additionally, the organoids were fixed to image fluorescence signal as index of mitochondrial activity by confocal imaging.

To investigate mitochondrial number, morphology, and size, VO samples were processed for transmission electron microscopy (TEM). VOs we’re fixed for 1 hour in 4% paraformaldehyde solution (Thermo Fisher Scientific, 7732-18-5) in PBS and for 2 hours in 2.5% glutaraldehyde in PHEM buffer (TAAB, G016). The PHEM buffer was composed of 5mM HEPES sodium salt (Sigma-Aldrich, H3784), 60mM PIPES sodium salt (Calbiochem, 6910), 10mM EGTA (EDM Millipore, 324626), 2mM magnesium chloride (Sigma-Aldrich, M4880). The Leica EMTP automatic tissue processor was used to process the VOs. The samples were washed in PHEM buffer and fixed in 1% osmium tetroxide (TAAB, O016). Serial dilutions of ethanol (Sigma-Aldrich, 51976) and Spurr low viscosity embedding kit (Electron Microscopy Sciences, 22050) was used to embed the VOs in resin. The prepared VOs were encased in Spurr resin and hardened at 60°C for 2 days and left to cool. The Leica Ultracut S UltraMicrotome was used to cut samples to 200 nm which were placed on Formvar-coated copper grids. Samples were stained with UA-Zero EM Stain (Agar, AGR1000) for 5 minutes, washed and stained with lead citrate (TAAB, L037) for 5 minutes. The JEOL JEM 1400plus TEM was used for imaging. Images were quantified using ImageJ using the Lam et al., protocol (Lam *et al*., 2021).

### 2.10. Generating HLI model, organoid labelling and injection, blood recovery assessment by Laser Doppler imaging, and cell tracking by Bruker imaging

To track and compare effects of human iPSCs-derived DB-VOs against ND-VOs on regeneration, 12 male NOD.CB17 Prkdcscid/NcrCrl mice (10-12 week-old) were randomly fall into two experimental groups of DB-VOs (n=6) or ND-VOs (n=6).The hindlimb ischemia model was created in these SCID mice by ligation of femoral artery (Margariti et al., 2012; Vilà-González et al., 2019). Before transplantation, equal number of DB-VOs and control ND-VOs were labelled by LuminiCell Tracker 670-Cell Labeling Kit (SCT011, Sigma-Aldrich), 0.8nm for 12 hours in Stempro34 complete medium. Then, 6-8 VOs (50 mg in total) per mouse were weighed and disrupted by 1 mL syringe (21 and 25 gauge) and the small clamps of cells in 200 µl PBS were injected intramuscularly into 3 points of the adductor muscle adjacent to site of ligation (the ischemic leg). Limb blood reperfusion was expressed as a ratio of the left (ischaemic) to right (non-ischaemic) paw at days 0, 1, and 14 post-operation, by Laser Doppler Perfusion System (Moor Instruments, UK).

At day 14, the mice were sacrificed and the muscles (skin off) of both legs were subjected to imaging with In-Vivo Xtreme Imaging System (Bruker, Germany). The biodistribution of labelled injected cells were spotted via Bruker Molecular Imaging (BMI) Software, and the area were harvested and embedded into plastic cylinder containing Optimal Cutting Temperature (OCT) compound and snap-frozen in precooled isopentane (2-Methylbutane) within liquid nitrogen. Again, the cylinders (on the dry ice within a petri dish) were imaged by Bruker system to exclude any cylinder without injected cells of VOs and to make sure where to exactly cryosection for subsequent histological analysis of injected cells and investigation of the integration into host vasculature.

### 2.11. Dissociation of the vascular organoids for single cell sequencing

To dissociate the VOs for single cell analysis, 5-10 VOs were transferred into 2 ml microtubes and washed 3 times with PBS. Then, they were mechanically disrupted by a 21G needle inside a 3 ml DLD dissociation solution (PluriSTEM® **D**ispase-II Solution, **L**iberase, #5401119001 **D**Nase I, #10104159001, all from Merck), and incubated for 20-30 min at 37°C. Cell suspensions were subsequently incubated in 50% Fetal Bovine Serum (Gibco) FBS-containing PBS on ice, filtered through cell strainers of 40µm and 30 µm, and centrifuged for 5 min at 500 g to get single cells pellets before staining with Flow Cytometry antibodies or proceeding to single-cell sequencing experiments.

### 2.12. scRNA-Seq Library preparation and sequencing

Twelve diabetic iPSC-derived VOs (n=3, 4 VOs per line) were evenly pooled together on day 21 of treatment, and the same procedure was performed in parallel for non-diabetic controls. They were dissociated within DLD solution and a 1 mL syringe (21 and 25 gauge), neutralized with 50% FBS in PBS, pelleted by centrifugation at 250*×g* and 4 °C for 5 minutes, washed once with 0.04% BSA in PBS, filtered twice with pluriSelect™ 40 µm/30 µm cell strainers. The viability of cells was double-checked both with trypan blue staining and automated counter Countess (Life Technology, AMQAX1000) to ensure >90% viability before sequencing. The single cells were diluted to 700-1200 cells/µL with 0.04% BSA in PBS and handed over to the Queen’s Genomics Core Technology Unit (GCTU) for immediate droplet generation (cell capturing) by 10X Genomics Chromium Controller. The Chromium Single-Cell 3′ Reagent Kits v2 Chemistry (10xGenomics) was used according to the manufacturer’s instructions. A total of 20,488 cells of dissociated VOs (7311 cells from a mixture of 3 non-diabetics, 13177 cells from a mixture of 3 diabetics) were captured for scRNA-seq library preparation and paired-end sequencing using the NovaSeq 6000 SP100 (Illumina) carried out at the GCTU of Queen’s University Belfast. Each sample provided ≈4 to 6×10^8^ 150 bp paired-end reads.

### 2.13. scRNA-Seq data processing

Reads’ mapping to human reference genome version 38 (GRCh38) and counting were processed using the Cell Ranger software v5.1.2 (10xGenomics). The output files containing the single cell counts were further analysed in the Partek Flow software (Partek Inc) and Bioconductor Packages in R. After the quality control step, the filtered expression matrix was normalized as count per million (CPM). Then, the principal component analysis (PCA) was performed on the counts of log2normalized Unique Molecular Identifiers (UMIs) to reduce the dimensionality of the features/genes. Using the first 20 principal components, t-distributed stochastic neighbour embedding (t-SNE) and uniform manifold approximation and projection (UMAP) were constructed on 2 or 3 dimensions to identify cell clusters. Subsequently, cell types were established based on the expression of their typical biomarkers.

### 2.14. Differentially expressed genes and Gene Set Enrichment Analysis (GSEA)

Differentially expressed genes were generated using the PartekFlow gene-specific analysis (GSA) package. Top statistically significant differentially expressed genes (DEGs) were obtained based on the criteria of FDR less than 0.05 and the absolute value of fold change between the two groups greater than 1.5. To exhibit the relative expression levels of the selected DEGs, the Enhanced Volcano plot, feature plots, and Boxplot/violin plot were generated using either PartekFlow or R 3.5.1. (Pheatmap package ver 1.0.12, ggplot2 packages ver 3.3.5). Normalized log2UMI expression values were used to produce the hierarchical clustering and Heatmaps. Pseudotime analysis was carried out using the Monocle2 package (ver 2.22.0) within PartekFlow.

Further downstream analysis was done by Gene Set Enrichment Analysis (GSEA). To take the gene variations into account, we calculated –log10 pValue of the DEGs and then multiplied it by the sign of fold change to get a metric. The ordered metrics were used to do pre-ranked GSEA using *fgsea* package *ver* 1.20.0 in R. Consequently, significantly positive and negative Hallmarks of the dataset for DB-VOs versus ND-VOs were represented on the GSEA plot.

## 3. Results

### 3.1. Successful generation of blood vessel organoids from iPS cells derived from diabetic and non-diabetic donors

An efficient and reproducible protocol for generating VOs is vital, to providing a promising drug-testing platform or to investigating underlying cellular and molecular characteristics in the *in vitro* model. In this study, colonies of iPSCs in 80-95% confluency **(Fig. 1a)**, were dissociated into cell clusters to induce the formation of homogeneous cell aggregates of 200 µm in size **(Fig. 1b)**. It was critical to not dissociate iPSCs into single cells but into 3-10 cell clusters for efficient aggregate formation (between 1000 to 1500 aggregates from one 6-well plate). The aggregates then underwent directed differentiation towards the mesodermal lineage and grew in form of floating spheroids **(Fig. 1c)**. Next, these floating spheroids were induced to initiate endothelial tube budding from the edge **(Fig. 1d)** to subsequently form vascular networks after embedding in the 3D Collagen I-Matrigel matrix **(Fig. 1e)**. Finally, every 3-5 vascular networks in the 3D environment were cut together on day 10 (**Supplementary Video 1);** allowing them to self-assemble into 3D floating, mature organoids culminating with smooth well-delimitated borders between days 15 to 20 **(Fig. 1f)**. The whole process from iPSCs to fully mature organoids is highly reproducible, and it is performed over a 20-day period. At that stage, the VOs (1-1.5 mm in size) were ready to be subjected to high glucose stimulation, drug screening, and/or any downstream analysis such as immunohistochemistry, protein, and RNA extraction for RT-PCR, WB, and protein arrays.

**Fig. 1.**
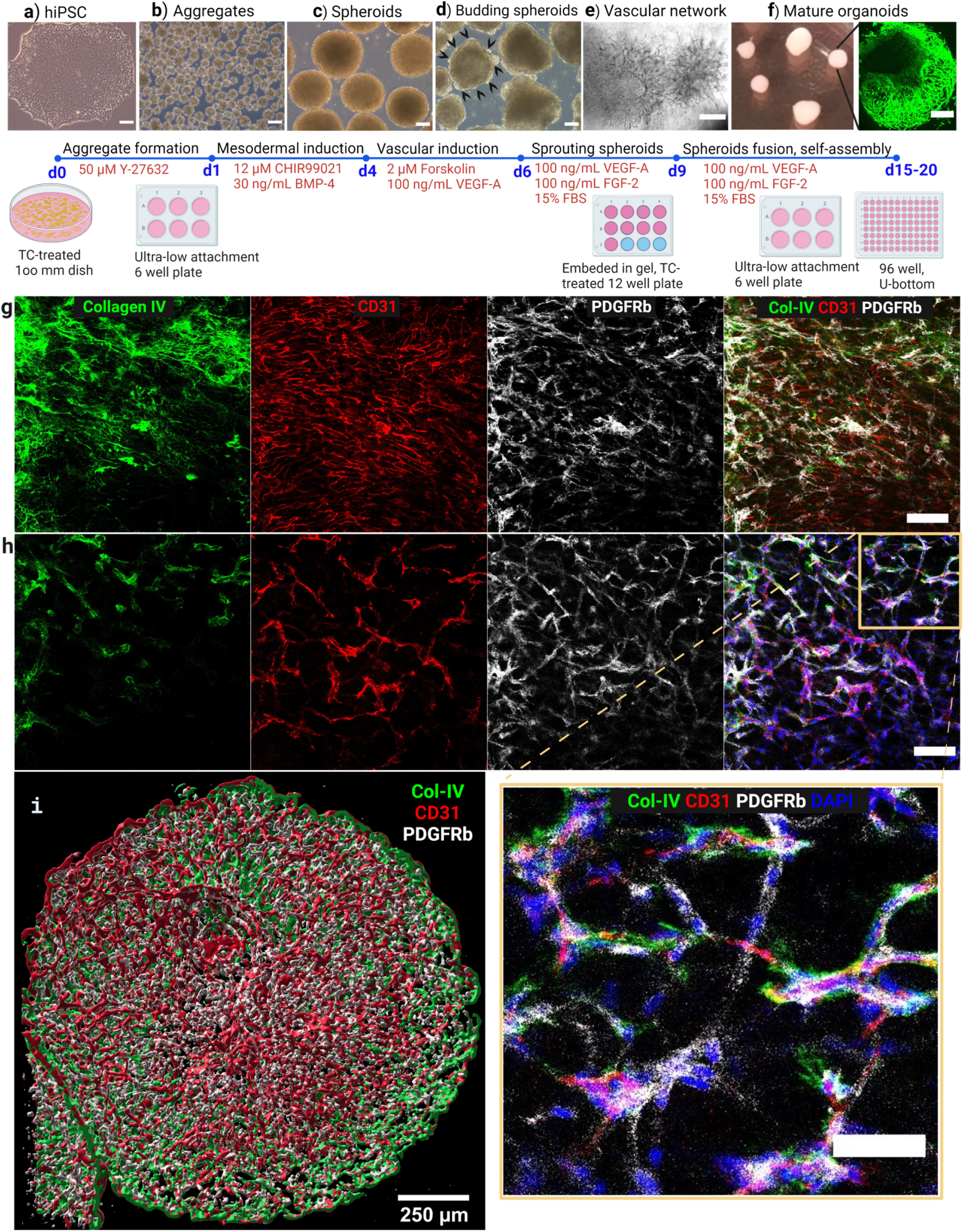
Blood vessel organoid generation and culture from iPSCs. **a**, Colony of human iPS cells in 80-90% confluency were dissociated for cell aggregate generation. **b**, Aggregate formation efficiency. The size of aggregates is on average 200 µm. **c**, Cell aggregates become bigger to form mesodermal floating spheroids. **d**, The floating spheroids start to bud after induction by VEGF-A and Forskolin. **e**, Phase-contrast image of the sprouting vessels that appeared 2 days after embedding in the 3D Collagen-Matrigel matrix. Every 2-4 vascular spheroids were cut to allow for making full, rounded, mature organoids till day 20 (**f**). Live staining of the mature organoids by Calcein-AM in green. Scale bars of A-F= 250 µm. Successful and reproducible generation of vascular organoids (VOs) from iPS cells that contain both small capillary networks (**g**) and bigger artery (**h**). Scale bar = 100µm. **Video** of the whole confocal series is represented in **Supplementary Materials**. The confocal imaging showed the presence of both endothelial tubes (CD31^+^, Red), mural cells (PDGFR-b^+^, Grey), and basement membrane (CollagenIV^+^, GREEN) within the VOs. The inset (Scale bar = 50µm) shows the interaction and alignment of ECs with mural cells. (**i**) 3D projection of 69 confocal images of a mature vascular organoid showing a nice alignment of mural cells with endothelial tubes. Scale bar = 200µm. Size range of organoids 0.5 to 1.5 mm.

To investigate cellular composition, the mature VOs were stained with antibodies against PECAM1 (CD31) and PDGFR-ß, to detect ECs and pericytes, respectively. Confocal images revealed the presence of both cell types **(Fig. 1g-i)**, confirming the successful generation of functional vascular networks from both diabetic and non-diabetic donors of iPSCs (**Extended Data Fig. 2**). As shown in **Extended Data Fig. 3a and b**, the primary plexus formation in VOs is mimicking the embryonic pattern of vascular development (Potente and Mäkinen, 2017). Interestingly, confocal images in the z-stacks showed the presence of both small network of capillaries **(Fig. 1g, Extended Data Fig. 4a and b, Supplementary Video 2)** and larger arteries as well **(Fig. 1h, Extended Data Fig. 4a and b, Supplementary Video 2)**, confirming full maturation and organization of the organoids. In particular, endothelial tubes in branching points were covered by pericytes as expected. Later, single cell sequencing data further confirmed the cell type composition and functions.

### 3.2. Diabetic vascular organoids revealed significantly enhanced ROS production and high Ac-LDL uptake by vascular cells

Early stages of atherosclerosis might be associated with elevated oxidative stress in the vasculature (Marchio *et al*., 2019), which can lead to vascular cells’ dysfunction and activate proatherogenic mechanisms. During environmental stress such as high glucose and proinflammatory cytokines in diabetes, ROS levels can increase dramatically leading to oxidative stress; this can result in damage to DNA, proteins, and lipids, and has been linked to CVDs (Luc *et al*., 2019). Accordingly, our results of confocal imaging (**Fig. 2a**) and Flow cytometry analysis (**Fig. 2b**) revealed significantly higher (median ratio=1.81, p value=0.001) ROS production in DB-VOs after 21 days of treatment with the diabetes-simulating stress (33 mM glucose, 1 ng/ml TNF-a, 1 ng/ml IL-6), as compared to ND-VOs.

**Fig. 2.**
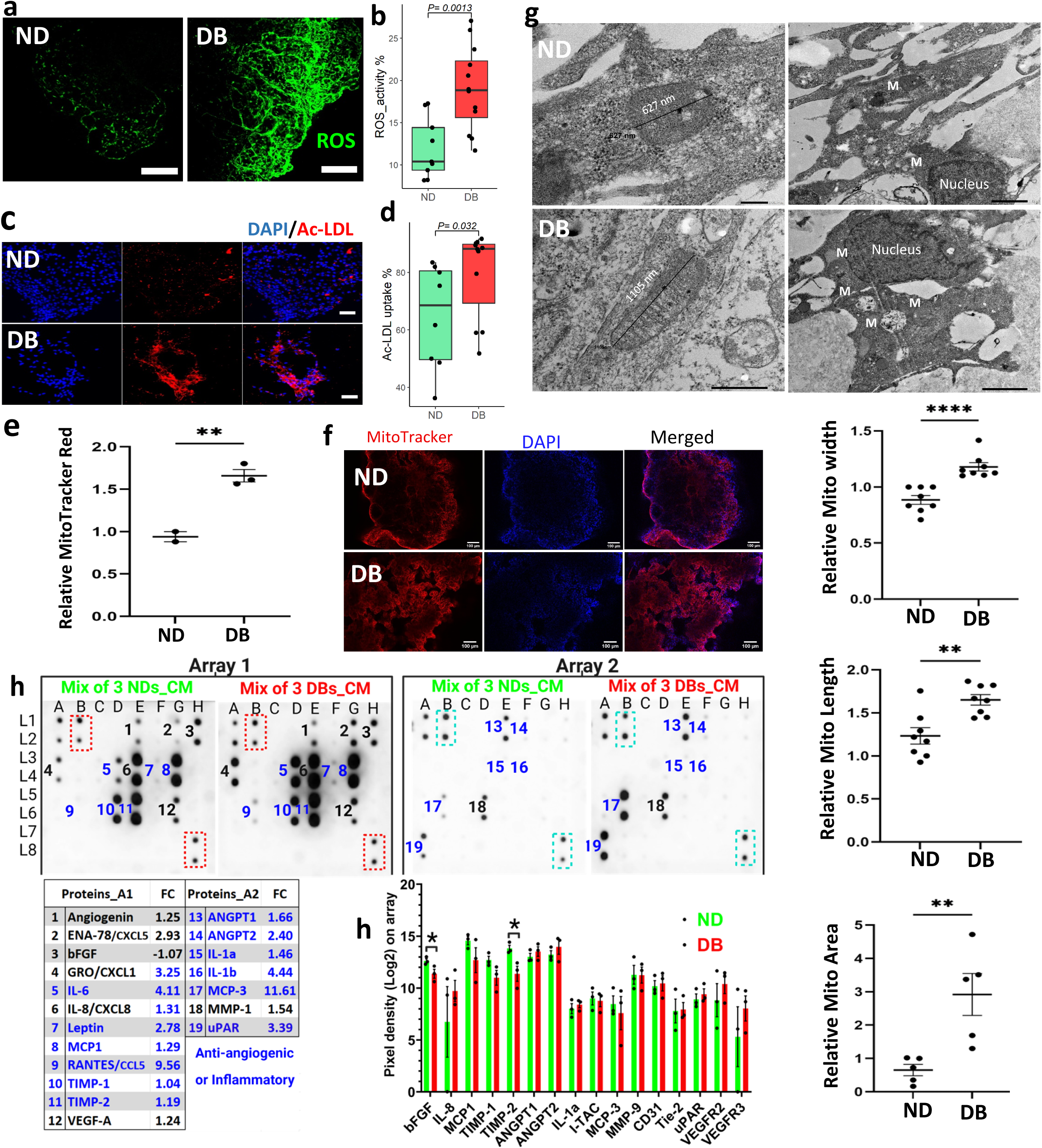
Functional assays of diabetic vascular organoids (DB-VOs) versus non-diabetic (ND-VOs). Enhanced ROS production in DB-VOs is represented by confocal imaging live (**a**), and a boxplot of flow cytometry analysis of 12 independent DB against 9 independent ND samples (**b**). Ac-LDL uptake showing by confocal imaging (**c**), and quantified by flow cytometry from 18 independent DB and ND samples represented by boxplot (**d**), which were significantly higher in DB-VOs as compared to ND-VOs. Scale bar in **a** and **c** is 100 µm and 200 µm, respectively. **e**, Quantification of MitoTracker Red Live Staining of vascular organoids in the bar chart showing significantly higher accumulation of mitochondria in DB-VO as compared to ND-VOs. **f**, Confocal imaging confirms the result of **e. g**, Transmission Electron Microscopy (TEM) shows larger mitochondria in DB-VOs as compared to ND-VOs. Error bars represent mean ± SEM (n = 3). The scale bars for ND are 200nm and 1µm, for DB 500nm and 2µm, left to right, respectively. P values are shown: **p < 0.01, ****p < 0.0001 (unpaired, two-tailed t-test). The human protein array of both conditioned media (minus FBS), 1.5 ml per well for 24 hours, the mixture of 3 DB-VOs versus 3 ND-VOs 6 weeks post-treatment (**h**), and 3 independent DB-VOs and ND-VOs individually 3 weeks post-treatment (**i**). 43 angiogenesis-related proteins were compared. 19 proteins were up/down-regulated in CM’s of DB at day 42 of diabetogenic media treatment, from which 12 of them were anti-angiogenic or pro/inflammatory cytokines (Shown in blue text on the arrays’ membrane and quantified in the table). DB VOs were treated with diabetogenic media (glocuse+TNFa+IL6) and control only with mannitol. Left panel: Array 1, and right panel: Array 2. CM; conditioned media. Values in the table represent the fold change (FC) ratio of each protein in DBs versus NDs. Dashed rectangles on the array membranes show the positive internal control for the normalization of each array. Bar plot in **i** showing the comparison of protein extracts from 3 independent DBs vascular organoids versus 3 independent NDs vascular organoids. Only bFGF and TIMP-2 were significantly different between DBs and NDs at 3 weeks post-treatment. A table showing the layout of all 43 antibodies against 20 proteins on Array 1, and 23 proteins on Array 2, along with the array membrane have shown in **Extended Data Fig. 6**. Unpaired t-test non-parametric (Mann-Whitney U test) was used to statistically examine differences between DBs and NDs. Mito or M denotes mitochondria.

Oxidative stress is central to endothelial dysfunction and it can be either a cause or a consequence (Odegaard *et al*., 2016). There is documented evidence that increased intracellular ROS primes vascular cells for the uptake of modified LDL, subsequently giving rise to the formation of vSMC-derived foam cells (Chellan *et al*., 2016). Elevated uptake of mod-LDL would elicit further vascular cells’ dysfunction, or activation, which participates in the initiation of atherosclerosis and exacerbates its progression (Xu *et al*., 2016; Tian *et al*., 2020). Studies have shown that uptake of modified LDLs (mod-LDLs) such as Ac-LDL (Jones, Reagan and Willingham, 2000), Ox-LDL (Tian *et al*., 2020), and glycated-LDL (Zimmermann *et al*., 2001), are the causes of foam cell formation, but not native LDL alone (Jones, Reagan and Willingham, 2000). However, there are cellular differences in the mechanism and the types of LDL receptors mediating the uptake of modLDL (Zimmermann *et al*., 2001; Veiraiah, 2005; Xu *et al*., 2016). Therefore, we decided to investigate differences in the uptake of Ac-LDL between DB and ND VOs. Interestingly, our results of confocal imaging (**Fig. 2c**) and Flow Cytometry (**Fig. 2d**) are showing that Ac-LDL is avidly taken up by vascular cells of the DB organoids as compared to ND organoids, with a median ratio fold of 1.28 (p value=0.03).

Live Staining of vascular organoids showed a significant accumulation of MitoTracker Red in DB-VOs as index of higher mitochondria activity (**Fig. 2e**), which further confirmed by confocal imaging (**Fig. 2f**). The number and size of mitochondria were significantly higher in DB-VOs as compared to ND-VOs (**Fig. 2g**). Transcriptome-based cellular component analysis of Gene Ontology (GO-CC) further confirmed the highly significant enrichment of mitochondrial compartments in DB-VOs versus ND-VOs (**Extended Data Fig. 5**).

### 3.3. Diabetic vascular organoids revealed significantly higher anti-angiogenic proteins in their secreted media

Cytokines are mainly produced by lymphocytes and macrophages, however, studies have shown that vascular cells are also implicated in inflammation and become dysfunctional after stimulation by injury or inflammatory mediators (Loppnow *et al*., 2011; Jha *et al*., 2018). Studies have also shown that transcriptional memory allows certain disease-related genes to respond more stronger to previously experienced signals, resulting in faster and greater signal-dependent transcription of the genes (Kamada *et al*., 2018; Zhao *et al*., 2020). Here, we simulated diabetes condition signals/inducers by a low-grade inflammation through adding 1 ng/ml of TNFa and IL-6, along with applying high glucose concentration (33 mM), to instigate pathological pathways in diabetic-patient iPSCs-derived VOs. Next, a human antibodies array (n=43) was used to compare secreted proteins on day 42 of treatment (**Fig. 2h and Extended Data Fig. 6a)** between ND-VOs and DB-VOs conditioned media as well as the protein extracts of the VOs on day 21 (**Fig. 2i and Extended Data Fig. 6b)**. The conditioned media collected from 3 independent biological replicates revealed enhanced secretion of anti-angiogenic proteins and inflammatory cytokines in the DB-VOs as compared to ND-VOs (blue text in **Fig. 2h**). Prominently, inflammatory cytokines RANTES, IL1b, and MCP3 had higher fold change of 9.56, 4.4, and 11.61, respectively, in secreted media of DB-VOs compared to ND-VOs at day 42 of treatment. However, comparing the protein extract of the DB-VOs (n=3) with the ND-VOs group (n=3) showed no significant difference between the two groups at day 21 of treatment (**Fig. 2i)**; Therefore, the higher expression of inflammatory cytokines at day 42 of treatment indicates that diabetic vasculopathy can worsen over time. This highlights the importance of time of incubation with pathology-simulating culture media to invoke epigenetic or metabolic memory of iPSC-derived vascular cells for greater transcription upon restimulation.

### 3.4. Diabetic vascular organoids expressed significantly less typical markers of ECs and mural cells

Analysing scRNA-seq data of vascular organoids on day 21 of treatment showed that DB-VOs as a whole have significantly lower expression of EC markers (PECAM1 and CDH5) and mural cells’ markers as compared to the ND-VOs (**Fig. 3a**). Applying a threshold of absolute fold change 1.5 and FDR=< 0.05, 680 genes were significantly upregulated and 1325 genes significantly downregulated, of which top 60 DEGs are shown in **Extended Data Fig. 7**. Functional enrichment analysis by GSEA based on the total DEGs of DB-VOs versus ND-VOs, revealed that Hallmarks of ROS and OXI-PHOS were significantly enriched in the DB-VOs (**Fig. 3b**), indicating pathology-simulating stress could significantly induce transcriptome changes that enrich for signalling associated with diabetes. Accordingly, the *in vivo* transplantation of VOs in an HLI recovery model showed that DB-VOs had 19.6% less blood recovery (**Fig. 3c**) at day 14 post-implantation (n=6, p value<0.01). Although the injected cells of both DB-VOs and ND-VOs were present at the sites of transplantation in the HLI region (**Fig. 3d**), confocal imaging revealed that the cells of human ND-VOs had increased integration with SCID mice vasculature as compared to DB-VOs. To understand which specific cell population is responsible for impaired function of DB-VOs; we further proceed with in depth analysis of single cell RNA-seq dataset.

**Fig. 3.**
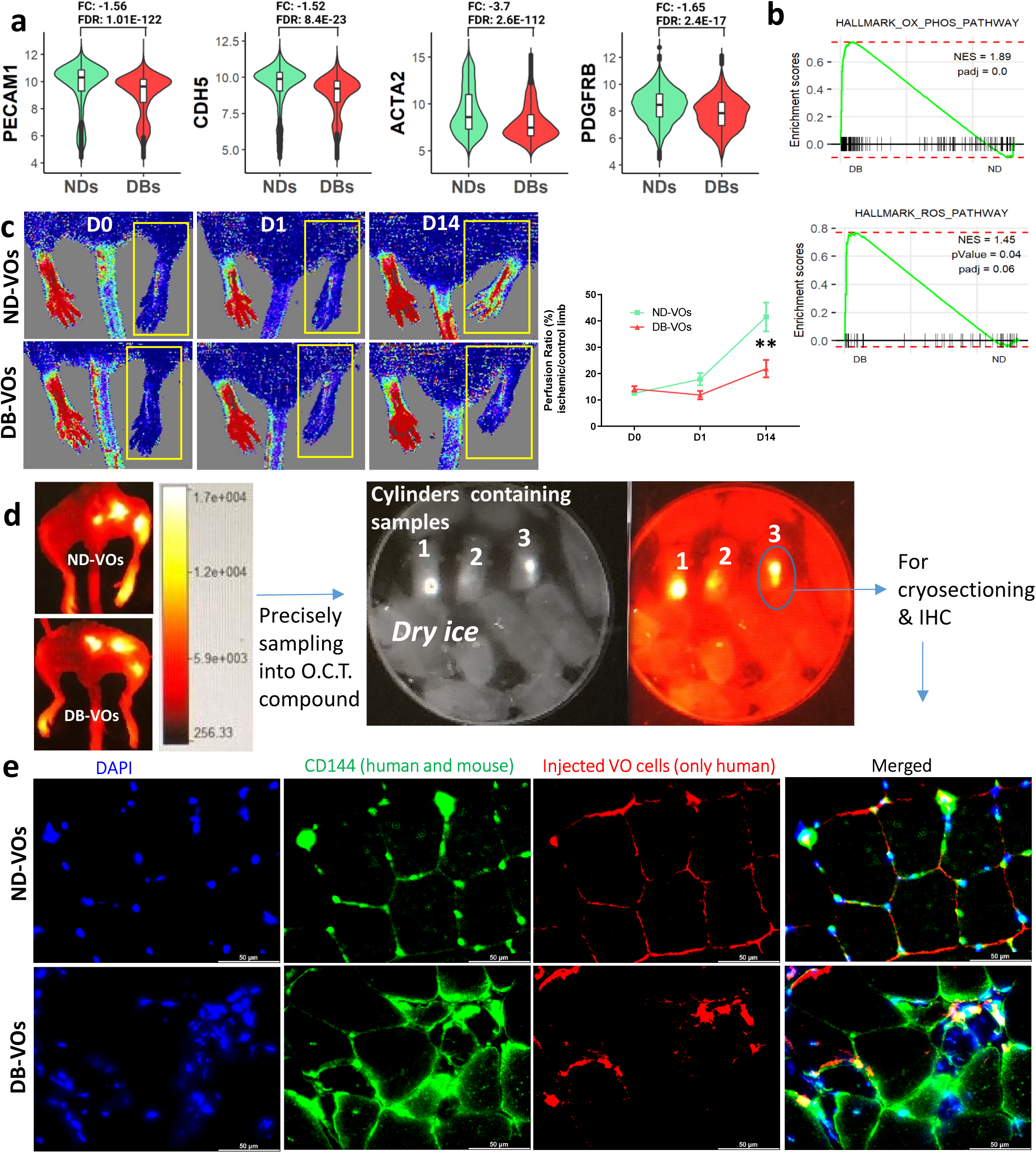
Comparison of diabetic vascular organoids (DB-VOs) with non-diabetic (ND-VOs) in terms of the transcriptome, Hind Limb Ischemia (HLI) recovery, regenerative potential, and integration into host vasculatures. **a**, Median value of log2-UMI (normalized expression value) is low for typical endothelial cell markers (CDH5 and PECAM1) in DB-VOs against ND-VOs as well as for mural markers (ACTA2 and PDGFRb). **b**, Based on the differentially expressed genes, DB-VOs are significantly enriched for the hallmark of ROS pathway and oxidative phosphorylation represented by GSEA plot. **c**, Laser Doppler imaging was performed on the HLI of the SCID mice on day 0 (immediately after surgery and transplantation of the VOs), day 1, and 14 post-surgery to assess blood re-perfusion in the lower limb, with the ischemic leg highlighted by the yellow rectangle, comparing the regenerative potential of injected DB-VOs versus ND-VOs, quantified in the line chart (n=6 per condition, statistical test two-way ANOVA, pvalue<0.01). The average blood perfusion recovery percent was 41.5% for ND-VOs against 21.8% for DB-VOs. **d**, The injected labelled cells around the HLI at 3 locations were tracked by Bruker imaging on day 14 after sacrificing mice. Then, the muscles were sampled precisely only from the injected area and embedded into Tissue-Tek O.C.T. Compound within cylinders. Again, the cylinders were tracked for labelled human cells by Bruker imaging on the dry ice to avoid heat damage to snap-frozen tissue samples. Only the cylinders that contain injected cells of labelled VOs further proceeded to cryo-sectioning. **e**, The sectioned slides were subsequently subjected to IHC and confocal imaging to see how the injected human cells of ND-VOs and DB-VOs were integrating into host vasculature. The injected cells of ND-VOs were better integrated with host vasculature.

### 3.5. Single-cell RNA sequencing data revealed specification of vascular cell types within iPSC-derived vascular organoids

To investigate cell composition and changes in ND-VOs and DB-VOs, the organoids from 3 independent iPSC lines for each group of DBs and NDs were pooled together and subjected to single cell transcriptome profiling on day 21 of treatment by 33 mM glucose, 1 ng/ml TNFa and IL-6. A total of 7,311 cells and 13,177 cells were captured in ND-VO and DB-VO samples, respectively (**Extended Data Fig. 8**). In the quality control step of single-cell data, cells with <500 detected genes as debris/low-quality cells (empty droplets containing apoptotic or dead cells’ particles) or cells with >7000 detected genes as the possible doublets/multiplets were excluded from subsequent analyses. Likewise, cells with low (<0.5%) or high (>50%) levels of mitochondria-encoded genes were filtered out (**Extended Data Fig. 8**). Therefore, totally 3782 cells out of 20,488 were excluded and the remaining 16706 VOs’ cells (5799 NDs and 10907 DBs sample), with ≈500-60,000 unique molecular identifier counts/cells were subjected to further downstream analysis. Dimensionality reduction was performed using principal components analysis (PCA) and the first 20 components were subjected to Graph-based unsupervised clustering and UMAP projection in 2 dimensions, resulting in 17 clusters (**Fig. 4a and 4b; Supplementary Excel Sheet 1**) in pooled DB-VOs and ND-VOs. Differentially expressed genes in DB-VO versus ND-VOs in each cluster are represented in **Supplementary Excel sheet 2**. The number of cells in each cluster varies, ranging from 151 to 2,623 cells per cluster (**Fig. 4b**). Additionally, we performed dimension reduction and Graph-based unsupervised clustering separately for ND-VOs and DB-VOs to interrogate diabetic-induced vascular cells’ heterogeneity based on transcriptome differences. The results indicated that DB-VOs have a more heterogeneous nature than ND-VOs, leading to 15 clusters (**Extended Data Fig. 9a and b**) against 12 clusters in ND-VOs (**Extended Data Fig. 10a and b**), owing perhaps to invoking inherent diabetic-related epigenetic or metabolic memory by the diabetogenic culture media. Each cluster possesses a unique set of differentially expressed genes (DEGs) as biomarkers (Macosko *et al*., 2015) that were used to annotate each cluster into a specific cell type (**Supplementary Excel Sheet 3 and 4)**.

**Fig. 4.**
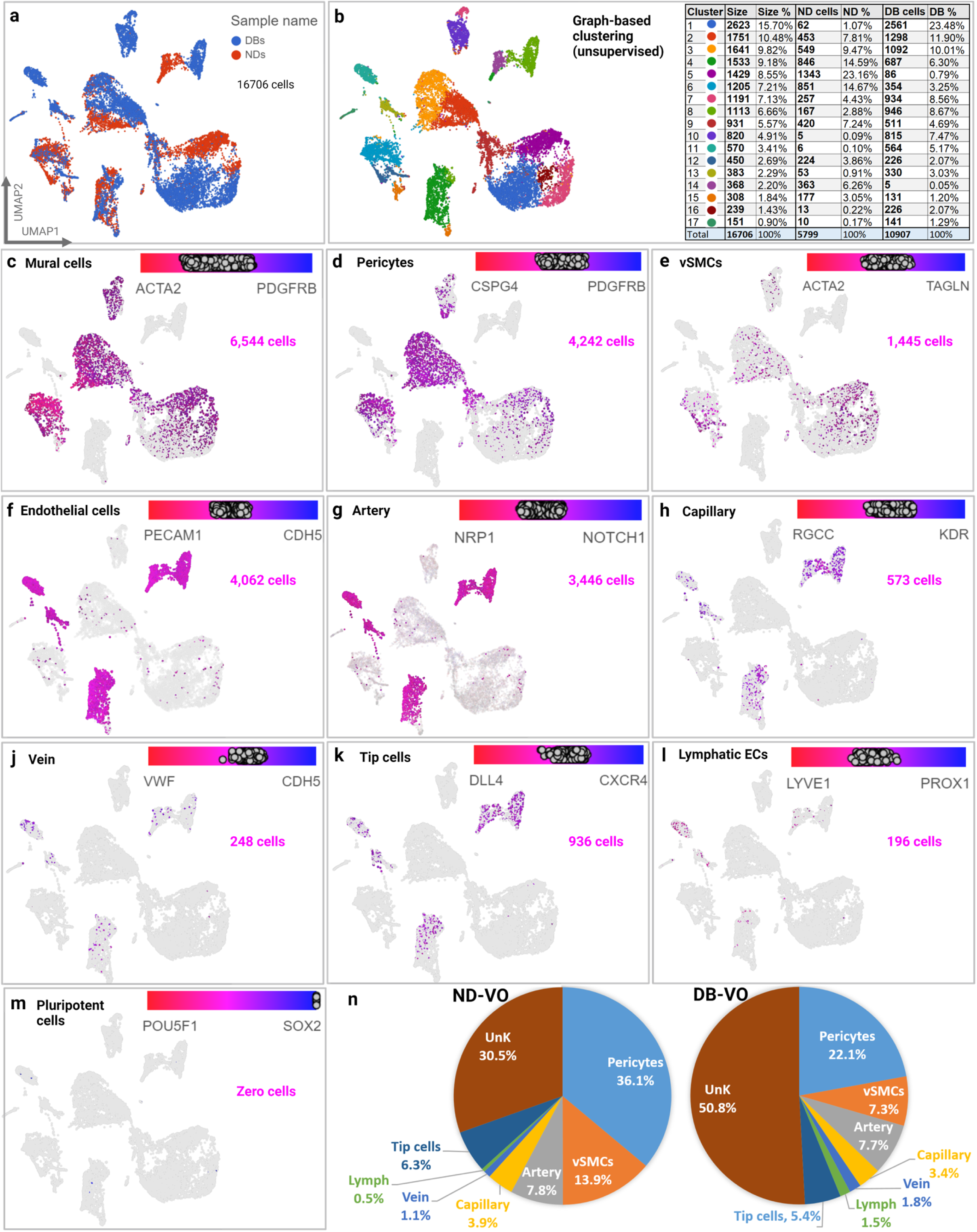
Vascular cell types’ specialization within blood vessel organoids (VOs). UMAP projection of the cells’ pool of VOs derived from diabetic (DB; 3 iPSCs lines) and non-DB (3 iPSCs lines) donors, coloured by the sample name in (**a**) and unsupervised graph-based clusters in (**b**). The number of clusters and their size is depicted in the table. One of the main populations within the VOs was mural cells (**c**) identified by PDGFRb and ACTA2 (a-SMA) markers, which were further classified into pericytes (**d**) expressing CSPG4/NG2 and PDGFRb and vSMCs (**e**) expressing ACTA2 and TAGLN. Additionally, endothelial cells, identified by PECAM1 (CD31) and CDH5 (CD144) markers, could further specialized into the artery (**g**) expressing NRP1 and NOTCH1, capillary (**h**) expressing RGCC and KDR, vein (**j**) expressing VWF and CDH5, tip cells (**k**) expressing DLL4 and CXCR4, and lymphatic ECs (**l**) expressing LYVE1 and PROX1. Of note, the cells expressing pluripotent stem cells’ markers were absent (**m**). (**n**) Piechart showing the percent of each cell type in ND-VOs versus DB-VOs.

Additionally, gene expression levels of traditionally well-known biomarkers were used to define clusters of mural cells (ACTA2+ and PDGFRb+) (**Fig. 4c)** and EC’ population (PECAM1/CD31+ and CDH5/CD144+) (**Fig. 4f)** on 2 dimensions of UMAP. We then further identified specification for vSMCs (ACTA2+ and TAGLN/SM22+) (**Fig. 4d)** and pericytes (PDGFRb+ and CSPG4/NG2+) (**Fig. 4e)**. Likewise, ECs could further be specialized into artery (NRP1+ and NOTCH1+), capillary (RGCC+ and KDR+), venous (VWF+ and CDH5+), tip cells (DLL4+ and CXCR4+), and lymphatic (LYVE1+ and PROX1+) (**Fig. 4g-l)**. Full list of biomarkers of each vascular cell type are presented in **Supplementary Excel Sheet 5**. Importantly, the cells expressing pluripotent stem cell markers (OCT4+ and SOX2+) were absent (**Fig. 4m)**, which would potentially give rise to teratoma with potential tumorigenic risk (Fu *et al*., 2012). Comparing the number of each cell type (**Fig. 4n)**, DB-VOs represented 3.1, 1.66, and 1.66 times higher lymphatic ECs, vein, and unknown cells, respectively, as compared to ND-VOs. On the other hand, populations of pericytes, vSMCs, artery, capillary, and tip cells were lower as compared to the ND-VOs (fold change ratios of 1.63, 1.89, 1.02, 1.17, and 1.17, respectively).

### 3.6. Single cell transcriptomic analysis revealed 2 distinct clusters of mural cells and ECs

By including a third dimension of the UMAP, two distinct far-distance clusters of cells were clearly seen within the pool of ND-VOs and DB-VOs (**Fig. 5a**). Further interrogation of these 2 clusters by Featureplot of classical mural cells-specific markers, ACTA2 and PDGFRb, and classical EC markers, PECAM1 (CD31) and CDH5 (CD144), confirmed that one of these clusters is a mural cell population (**Fig. 5b**) and the other one is ECs (**Fig. 5c**). The fact that the VOs contain two main populations was further confirmed by pseudotime and trajectory analysis in which a common parental cells (cell state 2) express mesenchymal stem cell markers, CD44, CD73/NT5E, CD90/THY1, and CD105/ENG. This common ancestor is further developed into a dichotomy; one branch point generates with the left-wing as mural cells (cell state 1) and the right-wing as ECs (cell state 3) (**Fig. 5d-g**).

**Fig. 5.**
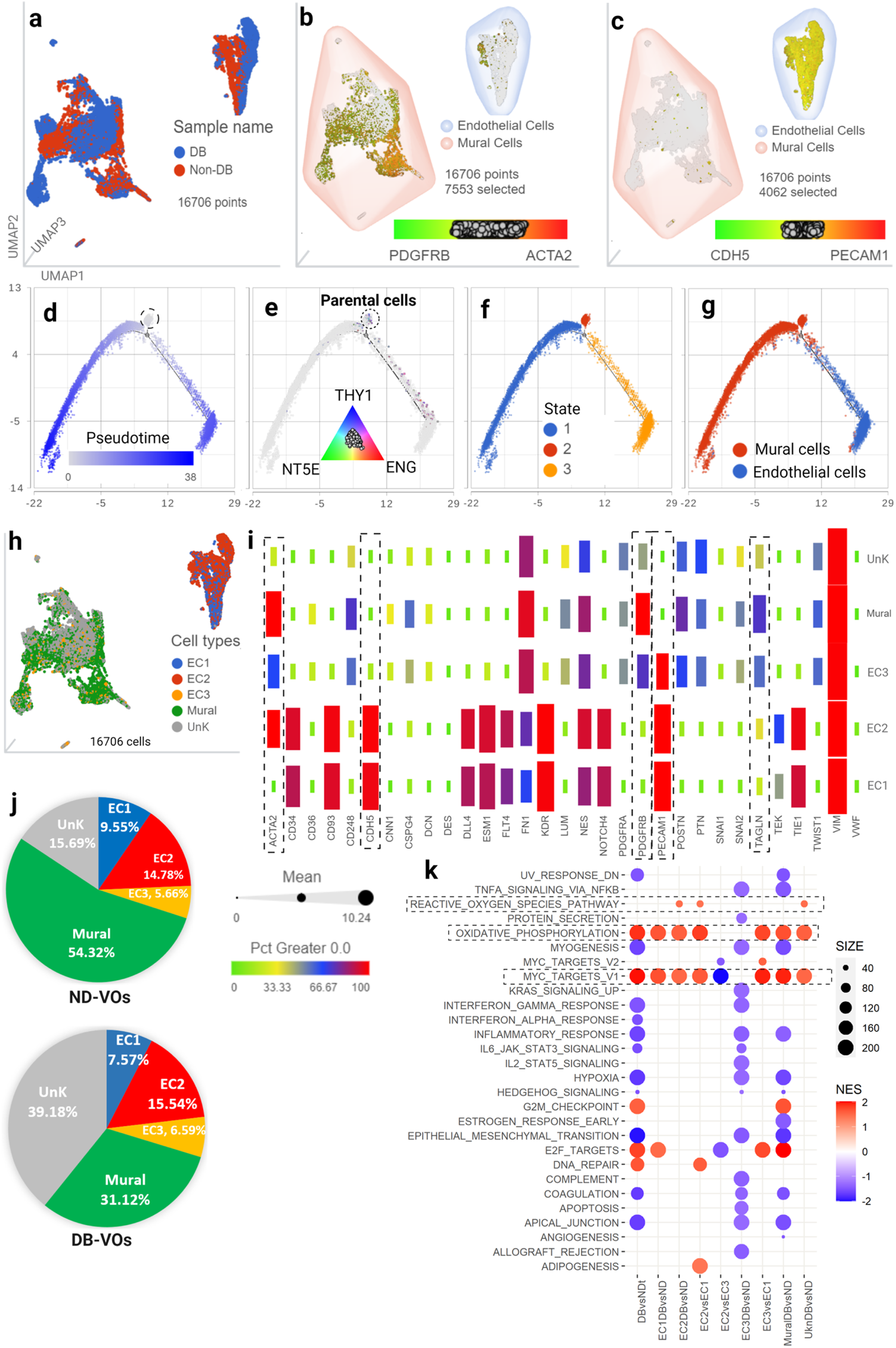
Dysfunctional EC subpopulation in diabetic vascular organoids (VOs) within the main EC cluster. 3D UMAP projection of the cells’ pool of VOs derived from diabetic (DB; 3 iPSCs lines) and non-DB (3 iPSCs lines) donors, coloured by the sample name in (**a**), mural cells’ subpopulation in (**b**) showing the expression of PDGFRb and ACTA2 (a-SMA), and ECs subpopulations in (**c**) showing the expression of PECAM1 (CD31) and CDH5 (CD144). **d and e**, Pseudo time analysis show a common ancestor cells expressing mesenchymal stem cell markers (CD44, CD73/NT5E, CD90/THY1, CD105/ENG), further developed into the more differentiated cell during the time (darker blue). **F**, VOs represent 3 cell states with a branch point, reflecting two differentiated cell populations with a common progenitor cells. **c**, overlying cell identities on the trajectory revealed that the dichotomy is mural cells in the left wing and ECs in the right-wing. **h**, After defining two main clusters of cell within VOs, we identified 3 subtypes of ECs within VOs’ cell pool on UMAP, named EC1, EC2, and EC3; all of which was expressing endothelial typical marker, CD31/PECAM1. However, EC1 and EC2 were within the EC cluster, but EC3 was within the mural cell cluster. **i**, A modified heatmap plot showing that EC2 within EC cluster is mesenchymal-like, expressing mesenchymal marker ACTA2 in addition to CD31 and CD144/CDH5. The level of expression (mean normalized log2UMI) has been represented by the size of the rectangles and the percent of cells expressing each marker has been shown by colour code from green, the lowest, to red, the highest. **j**, Piechart showing the percentage of each population in the ND-VOs and DB-VOs. **k**, Bubble plot showing Gene Set Enrichment Analysis (GSEA)-HALLMARKs based on differentially expressed genes of total cells of DB-VOs versus ND-VOs (DBvsNDt), EC1-DB versus EC1-ND (EC1DBvsND), EC2-DB versus EC2-ND (EC2DBvsND), EC2 versus EC1 (EC2vsEC1), EC2 versus EC3 (EC2vsEC3), EC3-DB versus EC3-ND (EC3DBvsND), EC2 versus EC3 (EC2vsEC3), mural cells-DB versus mural cells-ND (MuralDBvsND), and unknown cells-DB versus unknown cells-ND (UnkDBvsND). Only significant HALLMARKs at least at one condition are represented in the bubble plot. FDR <0.05 was considered significant.

### 3.7. A subpopulation of diabetic ECs represents impaired function with significantly higher hallmarks of oxidative phosphorylation and ROS pathways

Further interrogation of the two distinct clusters, revealed 3 subpopulations of ECs (**Fig. 5h**), named EC1, EC2, and EC3. EC1 was defined as a population of ECs that only express the EC markers within the ECs cluster, with the size of 9.55% in ND-VOs versus 7.57% in DB-VOs (**Fig. 5i**). Interestingly, there was also an EC2 population within the EC cluster that is expressing mesenchymal marker, ACTA2, with the size of 14.78% in ND-VOs versus 15.54% in DB-VOs (**Fig. 5j**). Of note, the EC2 population was highly expressing EC markers and ACTA2, but not TAGLN or PDGFRb (**Fig. 5i**), indicating that these cells are not possible doublets of ECs and mural cells. Additionally, there were some ECs within the mural cells’ cluster, named EC3, with the size of 5.66% in ND-VOs versus 6.59% in DB-VOs (**Fig. 5j**). In terms of cell percentage, there were fewer mural cells (31.12% versus 54.32%) and remarkably higher unknown cells (39.18% versus 15.69%) in DB-VOs versus ND-VOs (**Fig. 5j**). The unknown cells were defined as cells with no expression of typical markers of vascular cells. They were mainly located within the mural cell cluster but lacked expression of ACTA2. Biomarkers of these cell subtypes are represented in **Supplementary Excel Sheet 6**. Additionally, top cell-specific biomarkers of pooled cells from diabetic and non-diabetic vascular organoids have visualized in **Extended Data Fig. 11a**, along with the number of total counts and expressed genes, and the percent ribosomal reads and mitochondrial reads in each cell type and subpopulation for DB-VOs and ND-VOs, represented in Splitviolin plot (**Extended Data Fig. 11b)**.

To investigate what function is translated by the changes in the transcriptome of the DB-VOs population compared to ND-VOs, we performed GSEA (Hallmark) based on differentially expressed genes (**Supplementary Excel Sheet 7**). The significantly enriched hallmarks (FDR <0.05), either positive or negative phenotype in at least one of the conditions, are shown in the bubble plot (**Fig. 5k**). Of note, Hallmarks of oxidative phosphorylation and MYC-TARGETS_V1 are significantly (FDR <0.05) overrepresented in all conditions of diabetic versus non-diabetics. ROS is a by-product of dysfunction/electron leak from mitochondrial complexes 1-4 during oxidative phosphorylation across the respiratory chain (Shen, 2010). Comparing EC2 population versus EC1, hallmarks of oxidative phosphorylation (NES=1.81, padj=0.000) and ROS pathway (NES=1.43, padj=0.04) were significantly enriched in EC2 (**Fig. 6a**), based on differentially upregulated and downregulated genes (**Extended Data Fig. 12a**). The intersection of the ROS pathway gene set with the top significant DEGs list is represented in the bar chart, showing the FDR q-value and fold change of mean expression in EC2 versus EC1. EC2 population represents 3,803 biomarkers (**Fig. 6b, Supplementary Excel Sheet 8**), with the top 21 biomarkers shown in the inset table. Further interrogation of EC2-DB versus EC-2-ND revealed the enrichment of oxidative phosphorylation (NES=1.58, padj=0.000) and ROS pathway (NES=1.47, padj=0.003) are, indeed, originating from differentially expressed genes in the diabetic state (**Fig. 6c, Extended Data Fig. 12b, Supplementary Excel Sheet 7**). The intersection of the ROS pathway gene set with the top significant DEGs list is represented in the bar chart, showing the FDR q-value and fold change of mean expression in EC2-DB versus EC2-ND.

**Fig. 6.**
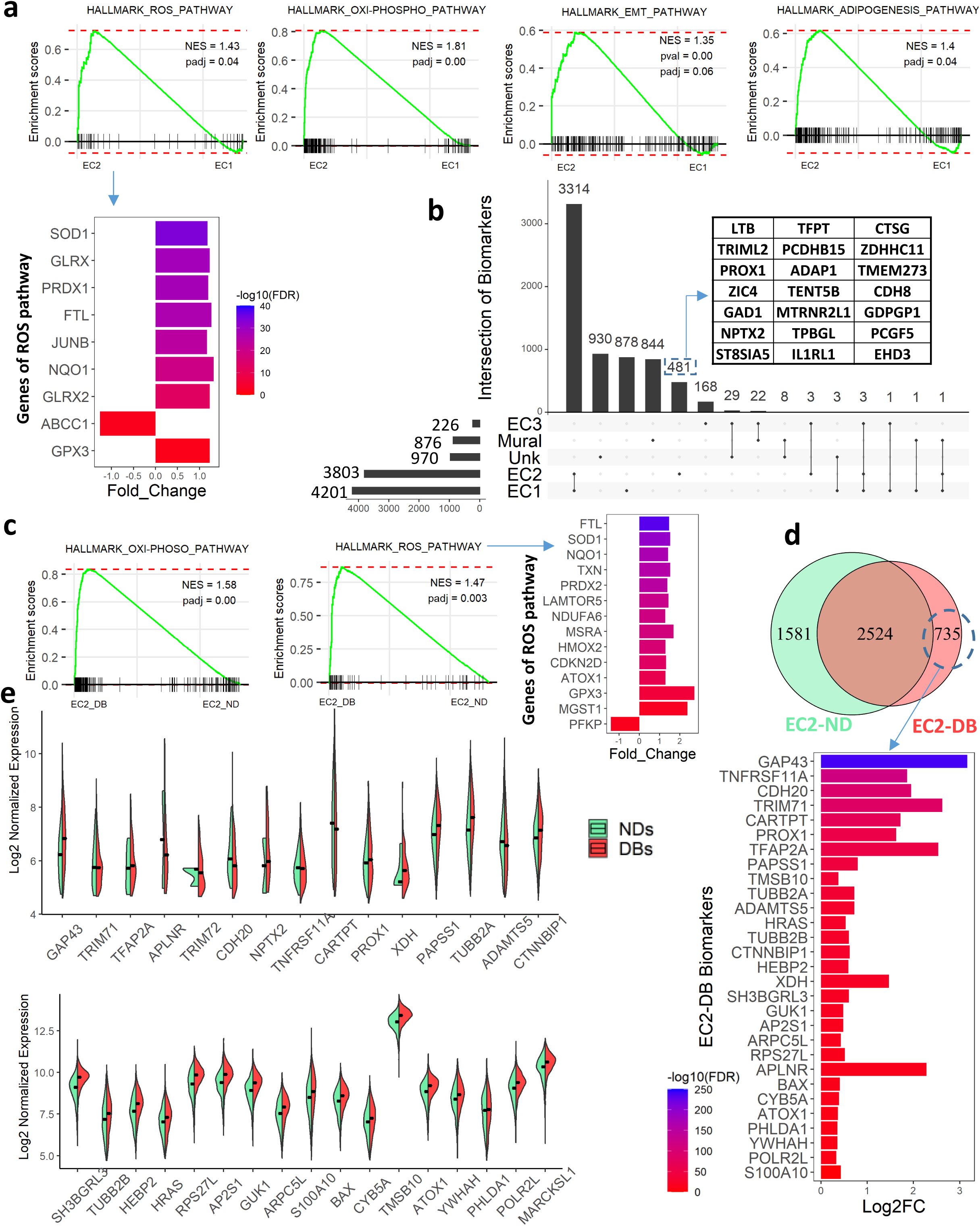
Biomarkers of EC2-DBs. **a**, Gene set enrichment analysis (GSEA) showing EC2 population overrepresents the hallmarks of adipogenesis, EMT, oxidative phosphorylation, and ROS pathway as compared to EC1, based on their differentially expressed genes (DEGs). The bar chart shows the fold change of the significantly expressed genes in EC2 annotated to the ROS pathway analysed by GSEA. **b**, UpSetR plot showing the intersections and unique biomarkers of each cell type within vascular organoids. The top 21 genes of EC2 unique biomarkers are shown in the table. **c**, Comparison of DEGs of EC2 population between DB-VO and ND-VOs showed significant enrichment for oxidative phosphorylation and ROS pathway in EC2 of DB. The bar chart shows the genes of ROS pathway that are significantly expressed in EC2-DB, which are responsible for the enrichment of the ROS hallmark. **d**, Venn diagram showing the common and unique biomarkers of EC2-ND and EC2-DB. The bar chart shows the top 30 genes of the intersection of the EC2-DB biomarker list (735 genes) with the list of significantly differentially expressed genes (DEGs; 7,611) in EC2-DB versus EC2-ND. **e**, Splitviolins showing median expression of top 30 DEGs of **d** in EC2 population. how to express in each cell type between DB-VO and ND-VO. Y-axis is the normalized expression value. The crossbar denotes the median expression for each gene.

EC2-DB specific biomarkers are potentially important to exclusively isolate these dysfunctional cells for further interrogation in the wet lab. Analysis of EC2-DB and EC2-ND biomarkers (**Supplementary Excel Sheet 9)** revealed 2,524 common biomarkers, 1581 EC2-ND unique biomarkers, and 735 EC2-DB unique biomarkers (**Fig. 6d**). The intersection of EC2-DB unique biomarkers with top significant DEGs is represented in the bar chart, showing the FDR q-value and log2 fold change of mean expression in EC2-DB versus EC2-ND. The median expression of these selected genes is represented by Splitviolin to show their changes in EC2 population DB versus ND (**Fig. 6e**).

## 4. Discussion

Vascular ECs are increasingly recognized as key players in CVDs. Yet, their functional properties and heterogeneity in diabetes remain uncharacterized. Studies have shown that ECs have basal heterogeneities in structure and function in different tissues and organs in the physiologic state (Gurevich *et al*., 2021; Pasut *et al*., 2021; Ricard *et al*., 2021); this makes therapeutic targeting of the endothelium challenging. However, for drug discovery, an understanding of the molecular regulators of EC function in the pathophysiological state is essential to be able to successfully interfere with the diabetic disease without affecting normal vasculature. Information about vascular cells’ subtypes mainly comes from studies of tissue or organ that contain many tissue/organ-imprinted confounding factors. Differences in tissue-specific features and functions of vascular cells, and existing other cells of microenvironment not relevant to disease make it even more complex by dynamically changing under the influence of many ill-defined processes (Langenkamp and Molema, 2009). Interestingly, diabetic iPSC-derived VOs have the advantage of not representing variations that arise from the tissue/organ-of-origin, while presenting heterogeneities originating from disease-related dysfunctionality of microvasculature with the advantage of having main vascular cells, without other confounding factors. Although there were differences between DB-VOs and ND-VOs in terms of the vascular cells’ percentage and ratio, we showed that VOs contain main vascular cells including different types of mural cells (pericytes and vSMCs) and ECs (artery, capillary, vein, lymphatic and tip cells). Therefore, iPSC-derived VOs can provide a suitable *in vitro* disease model, when induced by pathology simulating stressors, to unveil cellular and molecular characteristics/heterogeneities imprinted in the iPSCs’ memory of diabetic patients.

ECs perform an essential role in vascular function, however, their interaction with mural cells cannot be overlooked in how they respond to pathological conditions. Our results of confocal imaging and single cell data revealed that iPSC-derived VOs represent the benefit of having mural cells in addition to ECs, exhibited standard endothelial morphology and tube formation, nicely covered by pericytes. We were able to reproducibly grow full VOs within 15-20 days from iPSCs of both diabetic and non-diabetic donors, with an efficient differentiation protocol, unlike the previous protocol of 2D differentiation of only ECs from iPSCs (Paik *et al*., 2018).

As compared to control ND-VOs, DB-VOs showed significantly more ROS activity and modLDL uptake as well as higher pro-inflammatory cytokines over time, when challenged by diabetes-simulating stressors. These three early events (i.e. oxidative stress, modLDL uptake, and inflammatory cytokines) work in a vicious cycle in a way that one can worsen the other (Chellan *et al*., 2016; Odegaard *et al*., 2016; Jha *et al*., 2018; Marchio *et al*., 2019), would end up to diabetic vasculopathy and atherosclerosis.

We have used scRNA-seq as a profiling strategy to investigate the diabetes-specific features and intra-organoids mural cells and ECs heterogeneity in patient iPSCs-derived VOs at the single-cell level. We observed two distinct far-distance clusters expressing definitive biomarkers of either EC or mural cells, a dichotomy that was further confirmed by pseudotime trajectory analysis. We found that the most prominent cells within the VOs were arterial cells (3,446 out of 4,062 ECs) and pericytes (4,242 out of 6,544 mural cells). Within the main EC clusters, arterial endothelial genes were highly expressed, whereas venous endothelial genes had almost no expression, which was in agreement with (Zhang *et al*., 2017), even though Zhang *et al*. study was in iPSCs-ECs.

Our results showed that DB-VOs are more heterogeneous within mural cell cluster and possess 39.18% uncharacterized cells that do not express any typical markers of vascular cells. These findings are consistent with recent reports showing that vSMCs downregulate contractile markers during atherosclerogenesis and demonstrate multiple different characteristics by phenotypic switching to adopt alternative phenotypes (Grootaert and Bennett, 2021), indicating DB-VOs could recapitulate diabetic vasculopathy *in vitro*. Studies have also shown that vascular dysfunction is characterized by exaggerated proliferation of the vascular endothelium and/or smooth muscle cell contributing to the development of diabetic angiopathy and atherosclerotic plaque formation (Pinheiro-de-Sousa et al., 2022). These findings agree with our results showing a higher portion of proliferating unknown cells with enrichment of Hallmark MYC-TARGETS_V1 within DB-VOs. It has been reported that endothelial MYC overexpression results in severe defects in the human coronary artery (Pinheiro-de-Sousa *et al*., 2022) and the embryonic vascular system (Kokai *et al*., 2009). Similarly, high glucose has been shown to activate MYC target genes in different cell types (Jonas *et al*., 2001; Zhang *et al*., 2021), resulting in exaggerated cell proliferation. Of note, *Myc* is not only an oxidative stress-responsive gene (Elouil *et al*., 2005), but it can also induce ROS production (Vafa *et al*., 2002; Liu *et al*., 2017).

Although three EC subpopulations identified in this scRNA-seq profile are not limited to ND-VO but persist with DB-VO, only the EC2 population of diabetic vascular organoids represented impaired function (enriched for ROS activity pathway). This supports the idea that there is a “basal heterogeneity” from which different outcome arises depending on the transcriptomic memory and application of pathologic stressors (Kalluri *et al*., 2019). Previous studies have revealed that transcriptional memory allows certain disease-related genes to respond more stronger to previously experienced signals (Kamada *et al*., 2018; Zhao *et al*., 2020), similar to what we are observing in DB-VOs model.

The upregulation of contractile and migration gene (ACTA2/a-SMA) in the EC2 population in DB-VOs is particularly interesting given that endothelial to mesenchymal transition (EndMT) in atherosclerosis has previously been linked to EC dysfunction. (Chen *et al*., 2015; Kalluri *et al*., 2019). This was our rationale for focussing on the EC2 sub-population; however, there seem to be more gene differences between diabetic and non-diabetic organoids in the other subpopulations including uncharacterized cells. In agreement with our finding, a previous study have revealed that ACTA2-positive ECs only occasionally exist in thoracic aorta of wild type mice and rats, but frequently detected in 5-week-old apolipoprotein-E deficient mice. Interestingly, over time (in 20-to 24-week-old apolipoprotein-E deficient mice) the accumulation of these ACTA2-positive ECs dramatically increased especially in the luminal surface of atheromatous plaques, indicating the association of these EC2 subpopulation with progression of atherosclerosis (Azuma et al., 2009).

We found that DB-VOs had impaired mitochondrial function evidenced by changes in the number, size, and ROS production as compared to ND-VOs. In depth single cell analysis revealed that this impaired function is specific to the EC2-DB population, evidenced by a significantly higher normalized enrichment score for the ROS pathway (Hallmark) and mitochondrial cellular components (GO-CC). However, to confidently conclude that the EC2 sub-population is largely responsible for diabetic organoid dysfunction, our future efforts will aim to specifically separate and/or expand this subpopulation based on their unique biomarkers represented in this study. Besides the established vascular cell types within the vascular organoids, unknown or uncharacterized cells are also of great importance. Therefore, single-cell level transcriptomic profiling of DB-VOs versus ND-VOs in this study, provides an unprecedent potential for more discovery to find out what these yet unknown cells are and how they are functioning.

In conclusion, we reproducibly developed vascular organoids from iPSCs of both diabetic and non-diabetic donors, within 20 days, representing both main vascular cells. Enhanced ROS production and Ac-LDL uptake was the early signature of diabetic VO as compared to ND ones; confirming that lab-made VOs could recapitulate diabetic vasculopathy. Our study demonstrated that diabetic-related pathogenesis worsened over time, evidenced by enhanced anti-angiogenic proteins in conditioned media. Additionally, single cell RNA sequencing revealed less expression of endothelial and mural cells’ markers, more heterogeneity of cells, and more unknown cells in DB-VOs. Confirming the wet lab output, functional analysis of differentially expressed genes between three EC subpopulations in diabetic versus non-diabetic VOs showed significant enrichment of ROS and oxidative phosphorylation hallmarks in the EC2-DB versus EC2-ND; representing early signs of dysfunctionality in diabetic vascular organoids. The observed heterogeneity of ECs in the diabetic iPSCs-derived VOs suggests that diabetic disease-related responses might be specific to a small subpopulation of vascular cells. This study provides the list of unique biomarkers for subpopulations of DB-VOs, which is useful for further elucidation of molecular mechanisms of vascular diseases in diabetes.

## Supporting information

Supplementary Excel Sheet 1

Supplementary Excel Sheet 2

Supplementary Excel Sheet 3

Supplementary Excel Sheet 4

Supplementary Excel Sheet 5

Supplementary Excel Sheet 6

Supplementary Excel Sheet 7

Supplementary Excel Sheet 8

Supplementary Excel Sheet 9

## Author contributions

**Conceptualisation:** HN and AM. **Methodology:** HN, ME, GC, CN, AM. **Data collection and analysis:** HN, ME, CN, AM. **Data interpretation:** HN, ME, GC, VC, CN, SK, AY, CD, PDD, RA, AZ, LZ, AM. **Visualisation:** HN, ME, CN, RA, PDD, AM. **Writing original draft:** HN, AM. **Review and editing:** All authors. **Supervision and funding acquisition:** AM.

## Acknowledgments and Funding

This work was supported by grants from MRC (MR/X00533X/1), British Heart Foundation (PG/18/29/33731), and Northern Ireland Department for the Economy (USI-159).

The authors would like to thank Dr. Ileana Micu from Advanced Imaging, QUB, as well as staffs of Queen’s Genomics Core Technology Unit (GCTU) for technical expertise, support, and the use of instruments. We also thank Dr. Johnatas Dutra for helping with Bruker imaging.

## Data availability

All relevant data are within the paper and its Supporting Information files. The single-cell RNA sequencing data supporting the findings of this study is available from the corresponding author on reasonable request.

---

**Extended Data Fig. 1.**
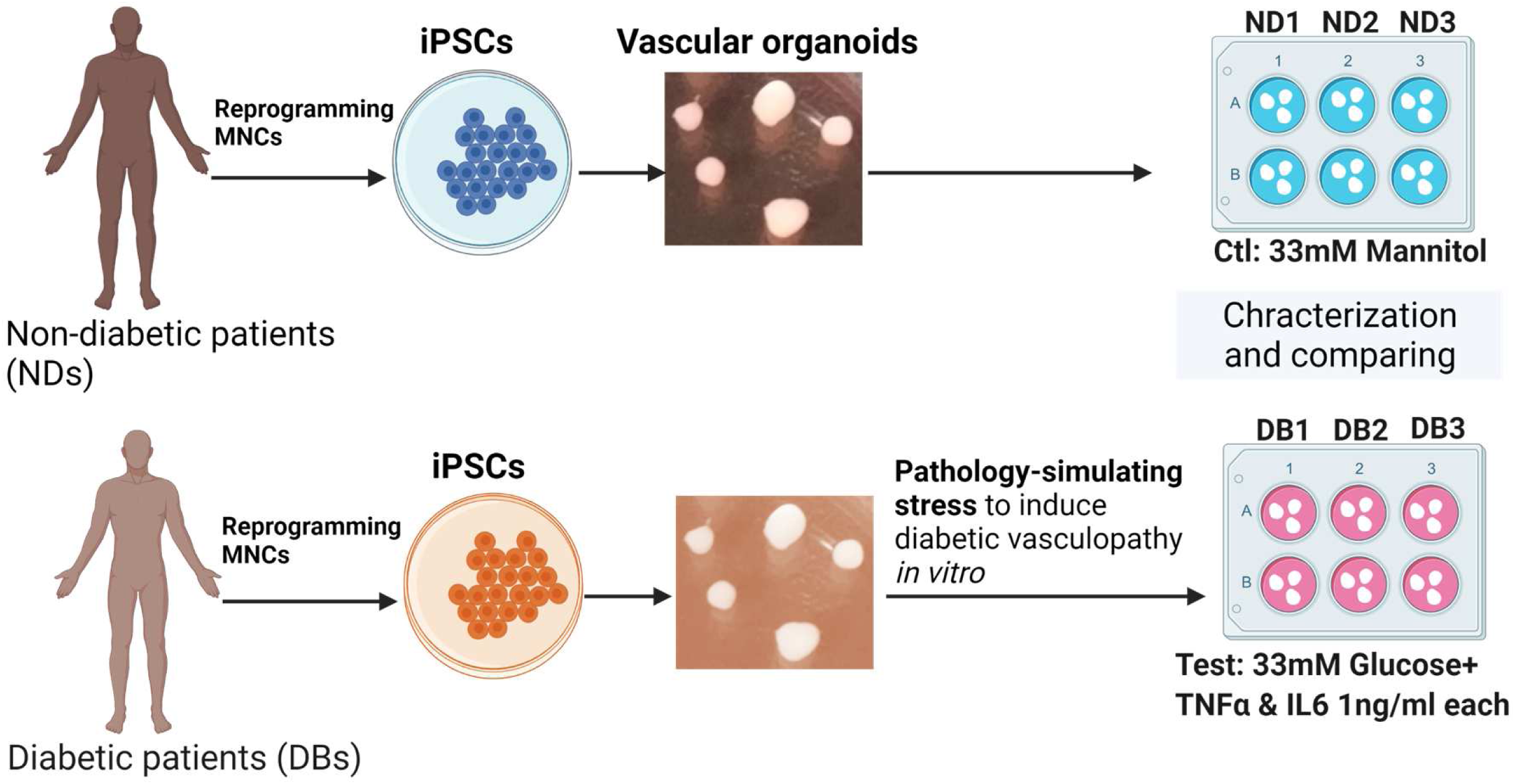
Experimental design to generate, characterize and compare vascular organoids from diabetic and non-diabetic individuals. iPSCs were reprogrammed from MNCs of NDs and DBs individuals and differentiated into vascular cells in a 3D manner to get diabetic and non-diabetic vascular organoids, called DB-VOs and ND-VOs respectively. To better disease modelling of diabetic vasculopathy/angiopathy *in vitro*, established mature DB-VOs at day 15 from diabetic iPSCs were started to culture in high glucose concentration (33mM) plus 1 ng/ml human TNF and 1 ng/ml IL-6 and compared to control ND-VOs within D-mannitol (33 mM), to maintain osmotic pressure the same as the test group. DB-VOs and ND-VOs were compared by both wet lab and dry lab analyses. For each analysis, at least three biological replicates were used.

**Extended Data Fig. 2.**
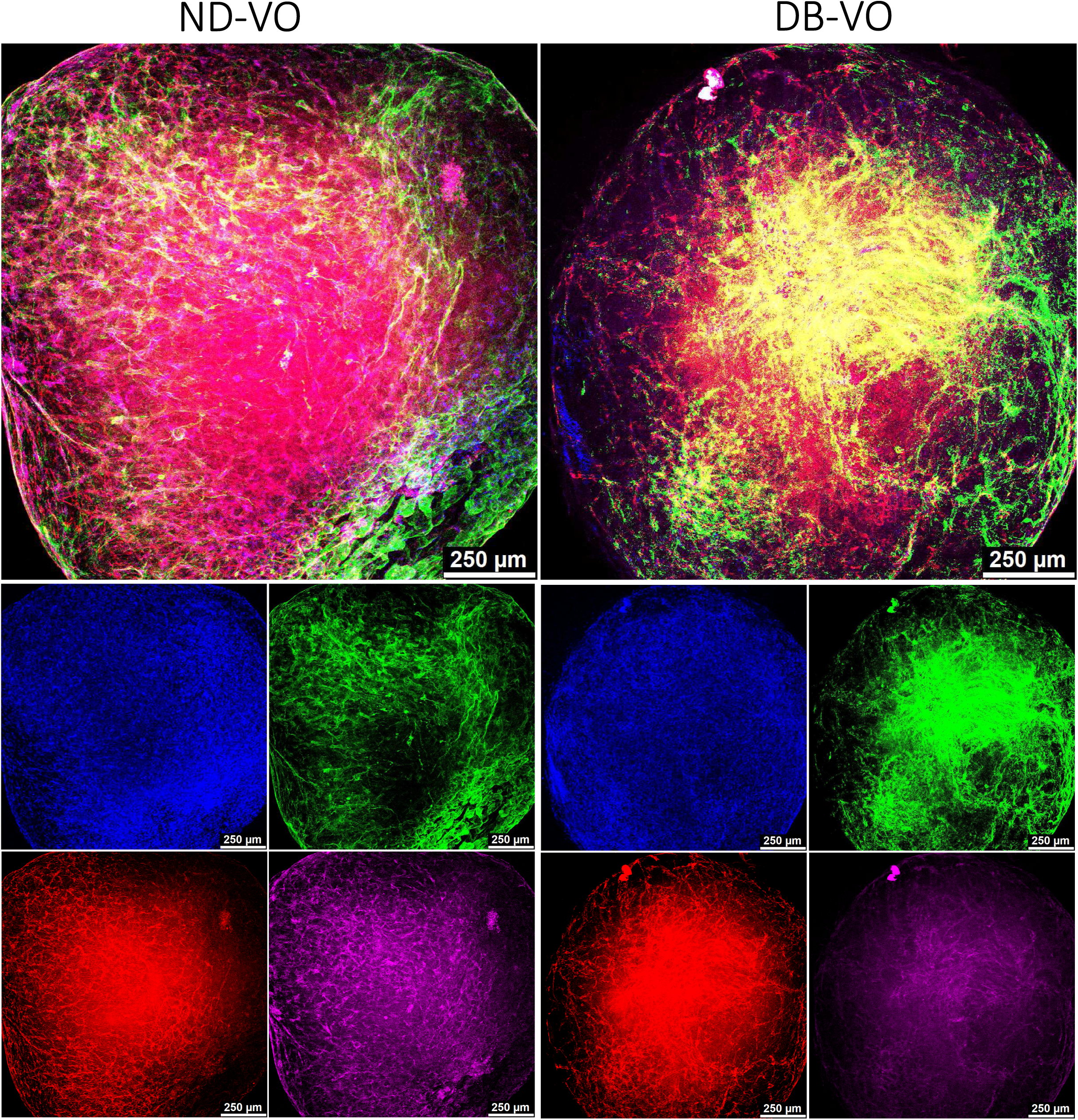
Generation of Vascular organoids from iPSCs of both diabetic and non-diabetic individuals. Successful and reproducible generation of vascular organoids (VOs) from iPSCs of both diabetic (DB) and non-DB (ND) individuals. Confocal imaging showed the presence of both endothelial tubes (CD31^+^, RED), mural cells (PDGFR-b^+^, MAGENDA), and basement membrane (CollagenIV^+^, GREEN) within the ND-VOs and DB-VOs. The upper panel is the merged image of its 4 separate channels in the lower panel. Blue is DAPI. Scale bar = 250µm. The size range of organoids was 0.5 to 1.5 mm.

**Extended Data Fig. 3.**
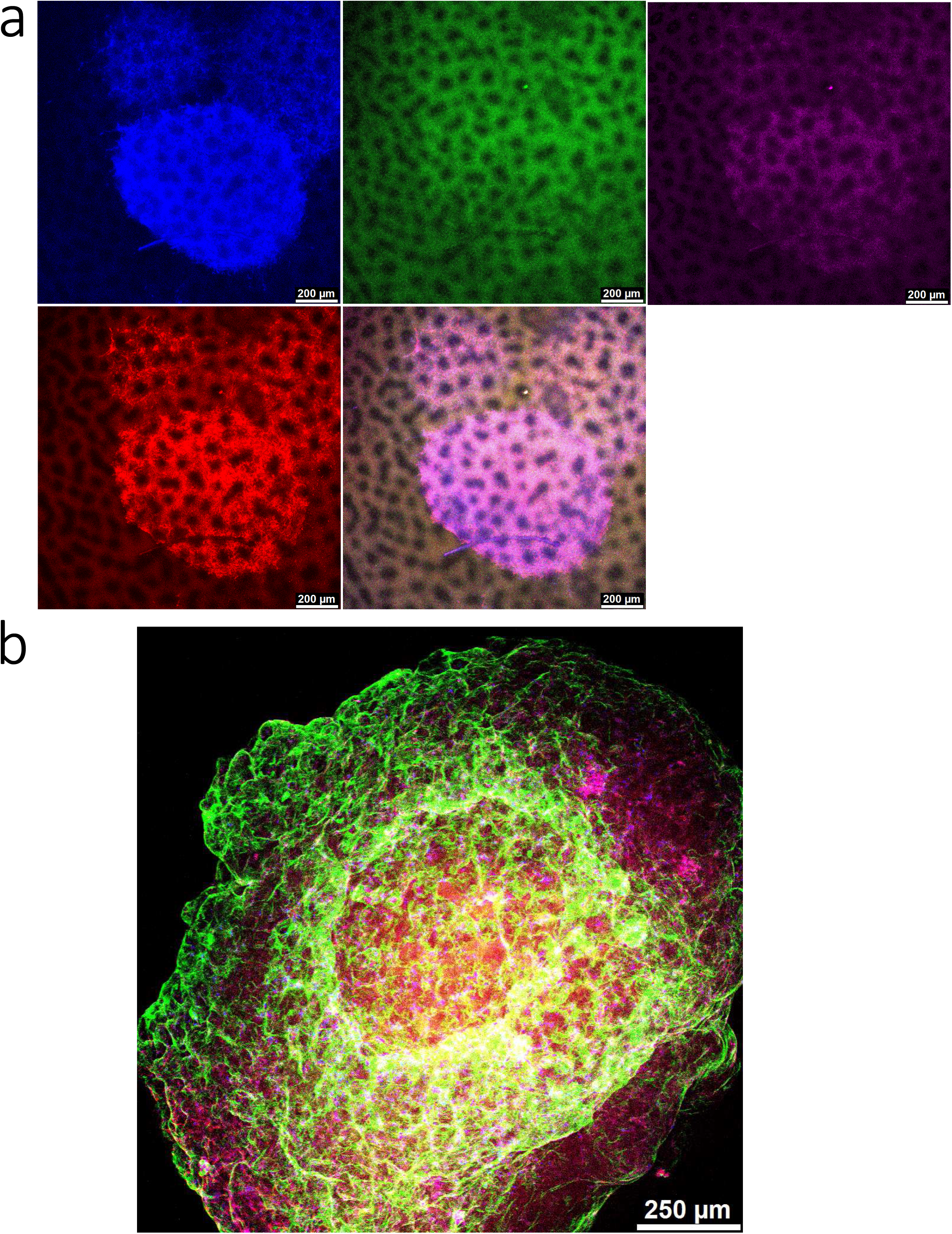
Generation of vascular organoids mimics the embryonic pattern of vascular development. Primary plexus formation in vascular organoids (**a**) that subsequently remodel into a network of capillaries, arteries, and veins (**b**). Endothelial tubes (CD31^+^, RED), mural cells (PDGFR-b^+^, MAGENDA), and basement membrane (CollagenIV^+^, GREEN).

**Extended Data Fig. 4.**
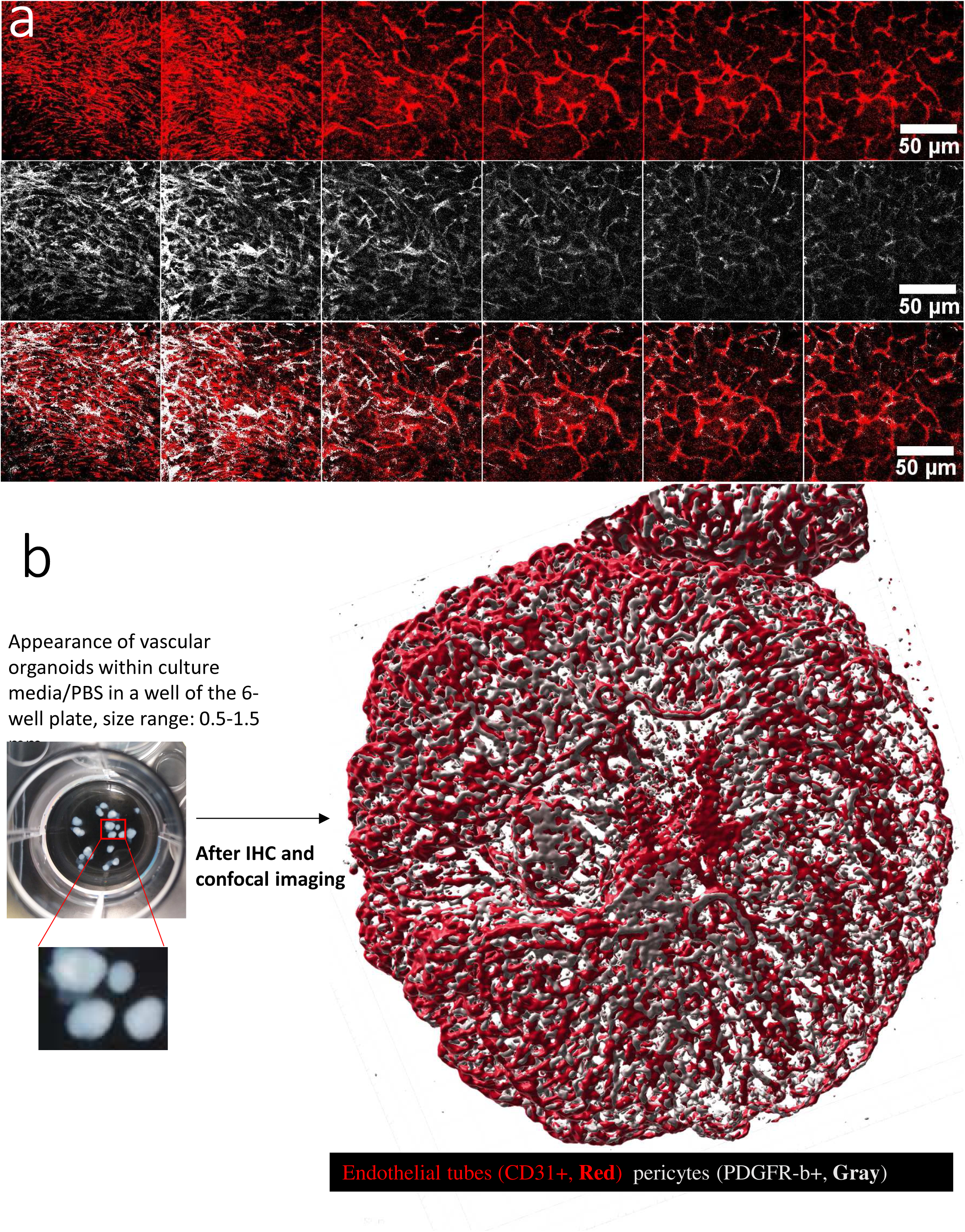
Presence of both small and large endothelial tubes. **a**, Slices of confocal images showing endothelial cells and mural cells and their alignment together within vascular organoids within the same field of Z-stacks. Endothelial tubes (CD31+, Red), mural cells (PDGFR-b+, Gray). **b**, Appearance of 3-dimensional vascular organoids in a tissue culture plate and after confocal imaging; showing how nicely pericytes are covering endothelial tubes.

**Extended Data Fig. 5.**
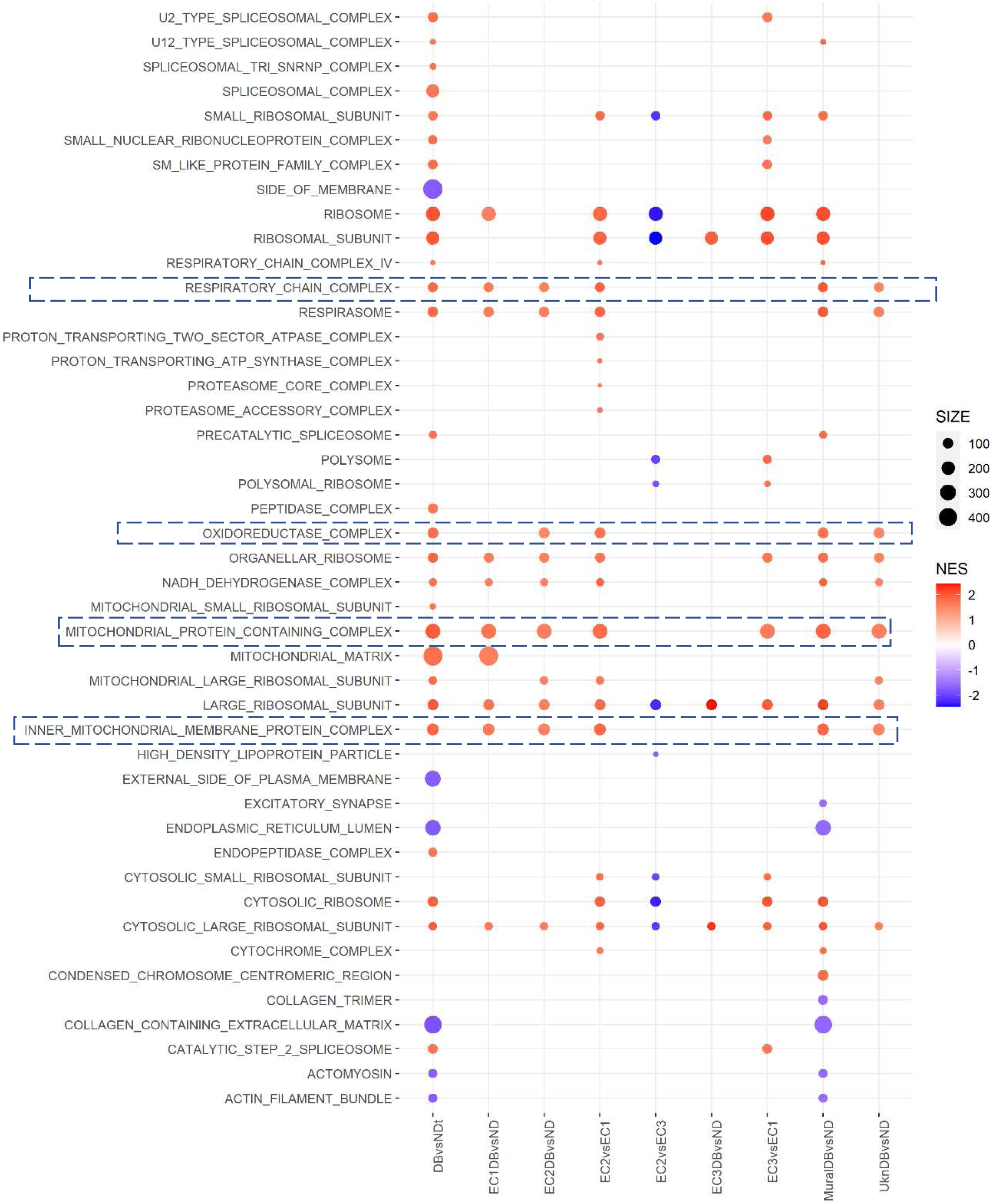
Enrichment of diabetic vascular organoids for mitochondria. A bubble plot of only significant (at least in one condition) Gene Ontology-Cellular Components (**GO-CC**) has been shown out of a total of 927 CCs. Mitochondrial components are highly enriched in DB-VOs versus ND-VOs. FDR <0.01 was considered significant.

**Extended Data Fig. 6.**
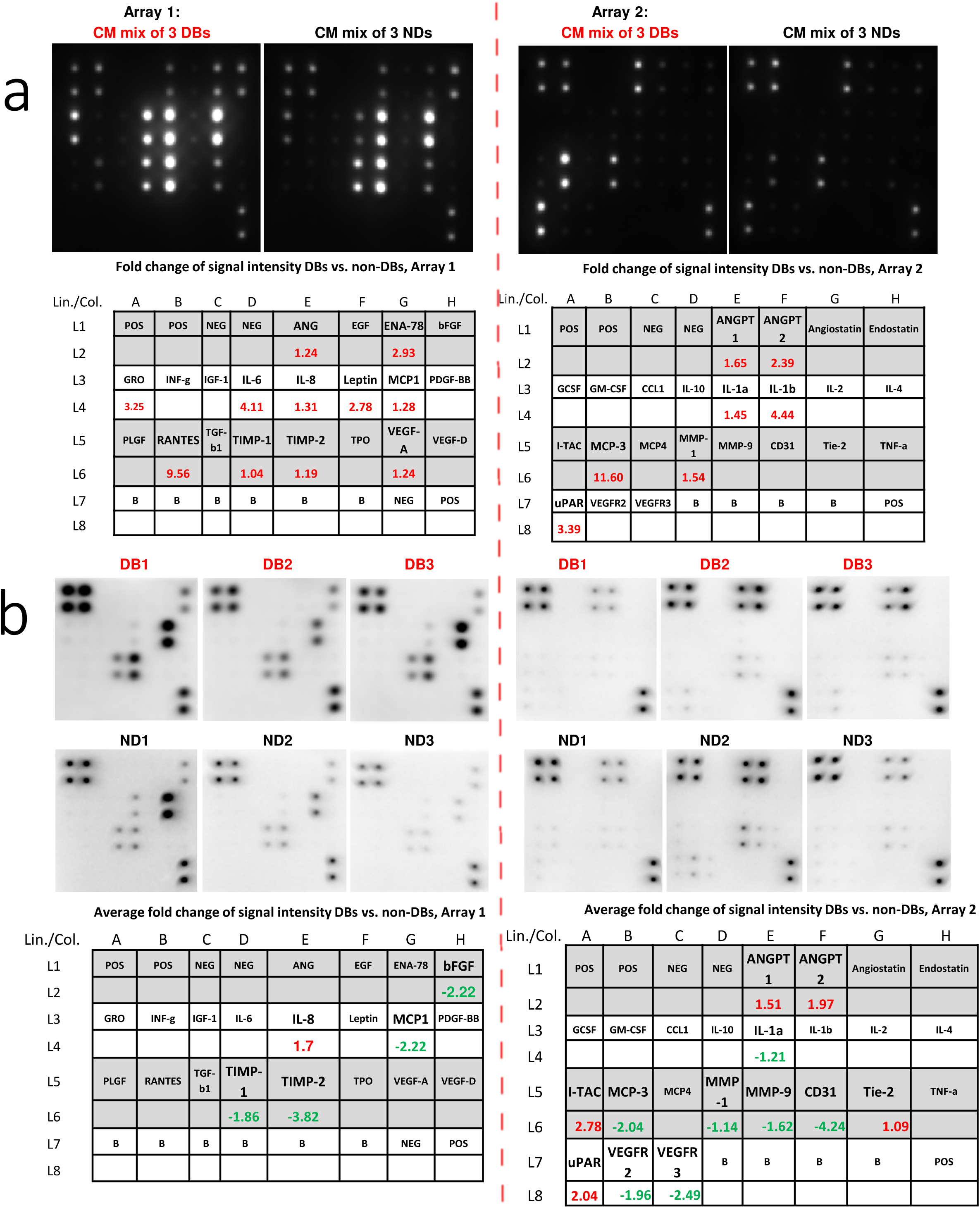
Human protein array of diabetic and non-diabetic vascular organoids. **a**, Comparison of the conditioned media (minus FBS), 1.5 ml per well for 24 hours, the mixture of 3 DBs versus 3 NDs vascular organoids. DB-VOs were treated with g+TNFa+IL6 and control only with mannitol. **b**, Comparison of protein extracts from 3 independent diabetic vascular organoids versus 3 independent ND-VOs. Left panel: Array 1, and right panel: Array 2. Tables showing the layout of antibodies against 20 proteins on Array 1, and 23 proteins on Array 2, with values representing the fold change ratio of each protein in DBs versus NDs.

**Extended Data Fig. 7.**
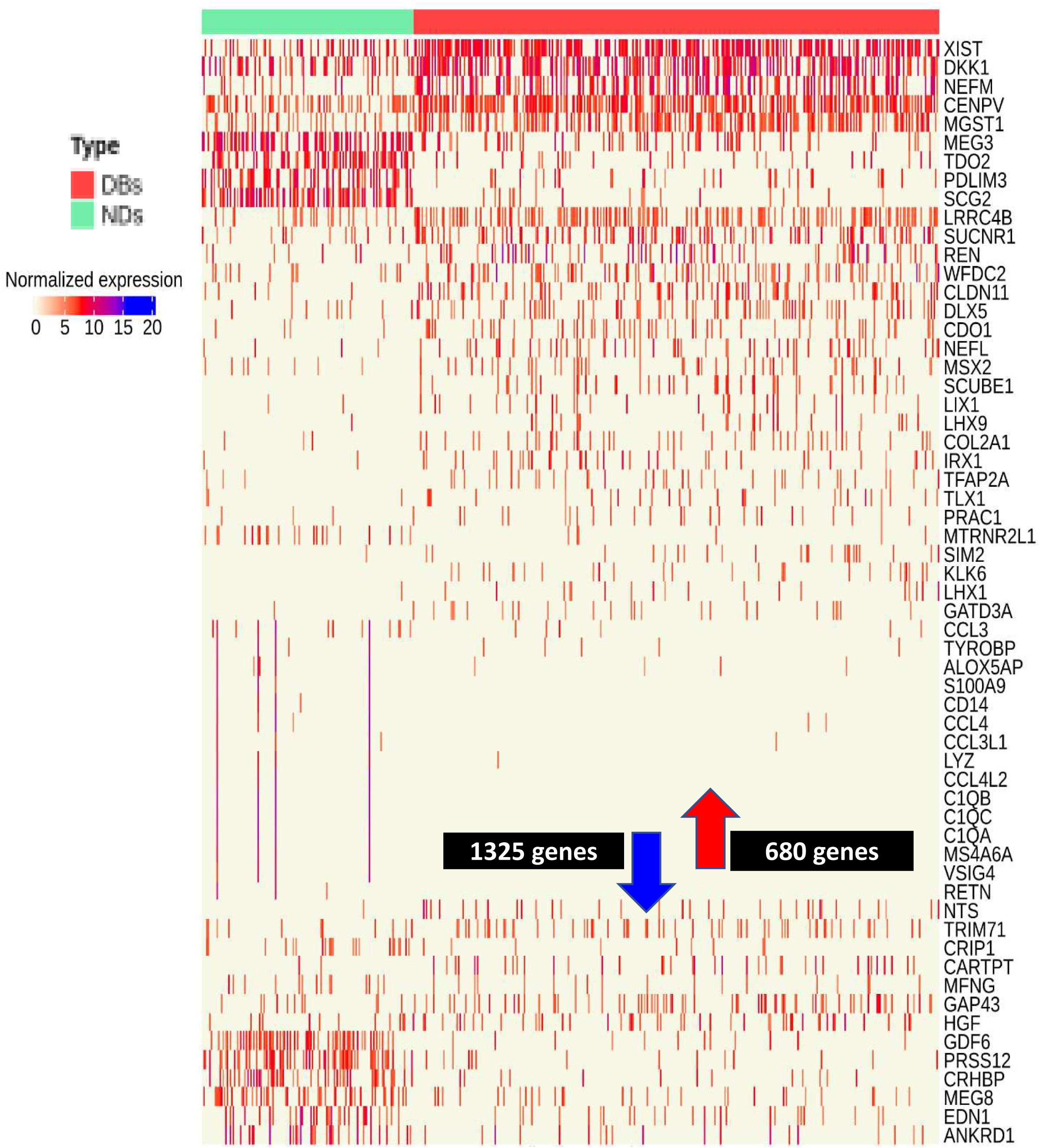
Differences of normalized Log2-UMI expression values for top 60 differentially expressed genes (DEGs) in DB-VOs versus ND-VOs. In total, 680 genes were significantly upregulated and 1325 genes were significantly downregulated in DB-VOs. Genes with an absolute fold change of 1.5 and FDR<0.05 were considered significant changes. The list of DEGs was used for downstream functional analyses. UMI: Unique Molecular Identifier.

**Extended Data Fig. 8.**
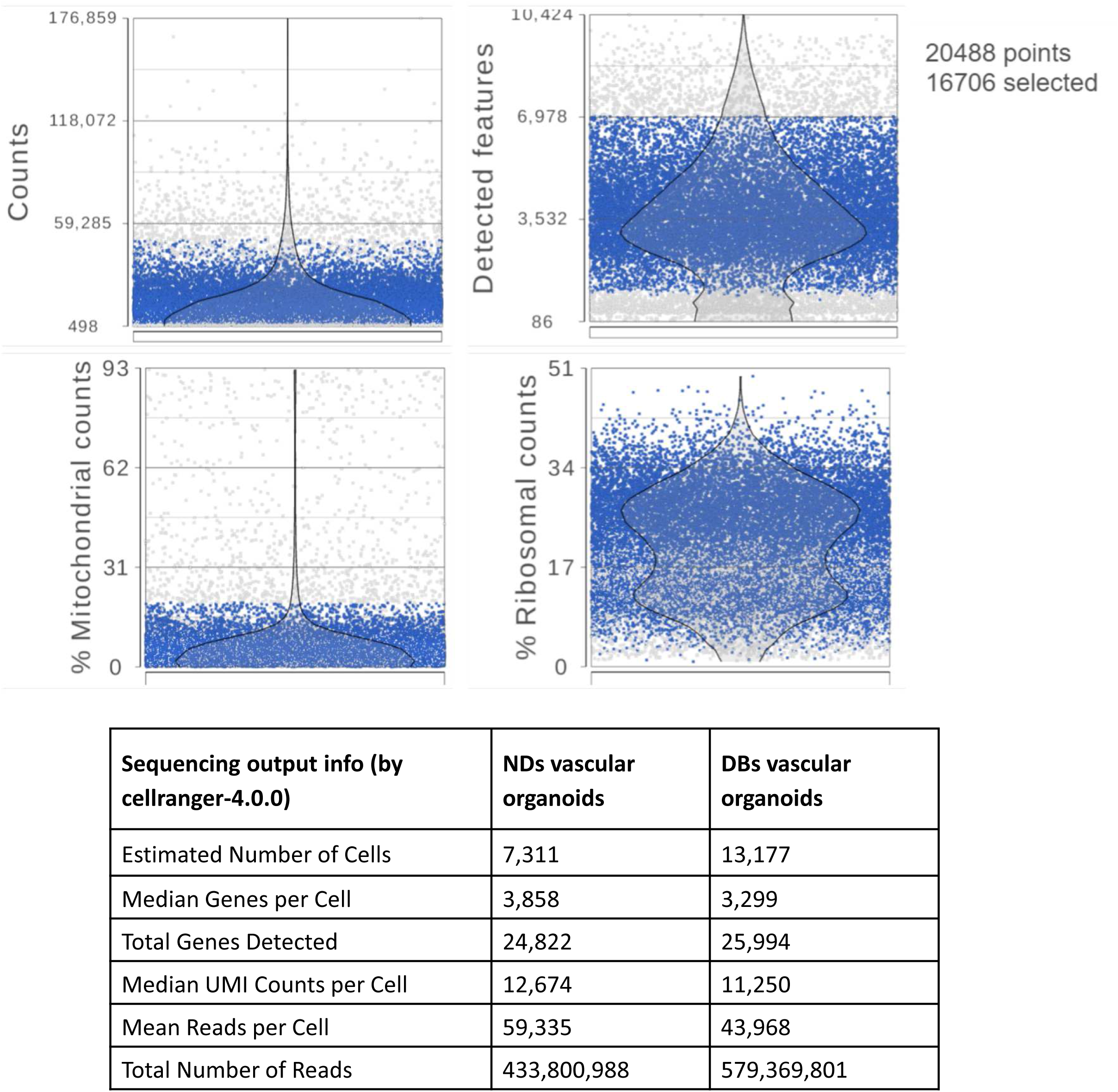
Quality control trimming of single cell data. In the quality control step of single-cell data, cells with >7000 detected features (genes)/cell as the possible doublet/multiplet cells were excluded. Additionally, cells with <500 detected feature/cell and with low (<0.5%) or high (>50%) mitochondrial read counts as debris/low-quality cells (empty droplets containing apoptotic or dead cells’ particles) filtered out. As a result, the remaining 16706 vascular organoids’ cells (5799 ND and 10907 DB cells) out of a total of 20,488 cells), with a range of ≈500-60,000 unique molecular identifier counts/cell, were subjected to further downstream analysis.

**Extended Data Fig. 9.**
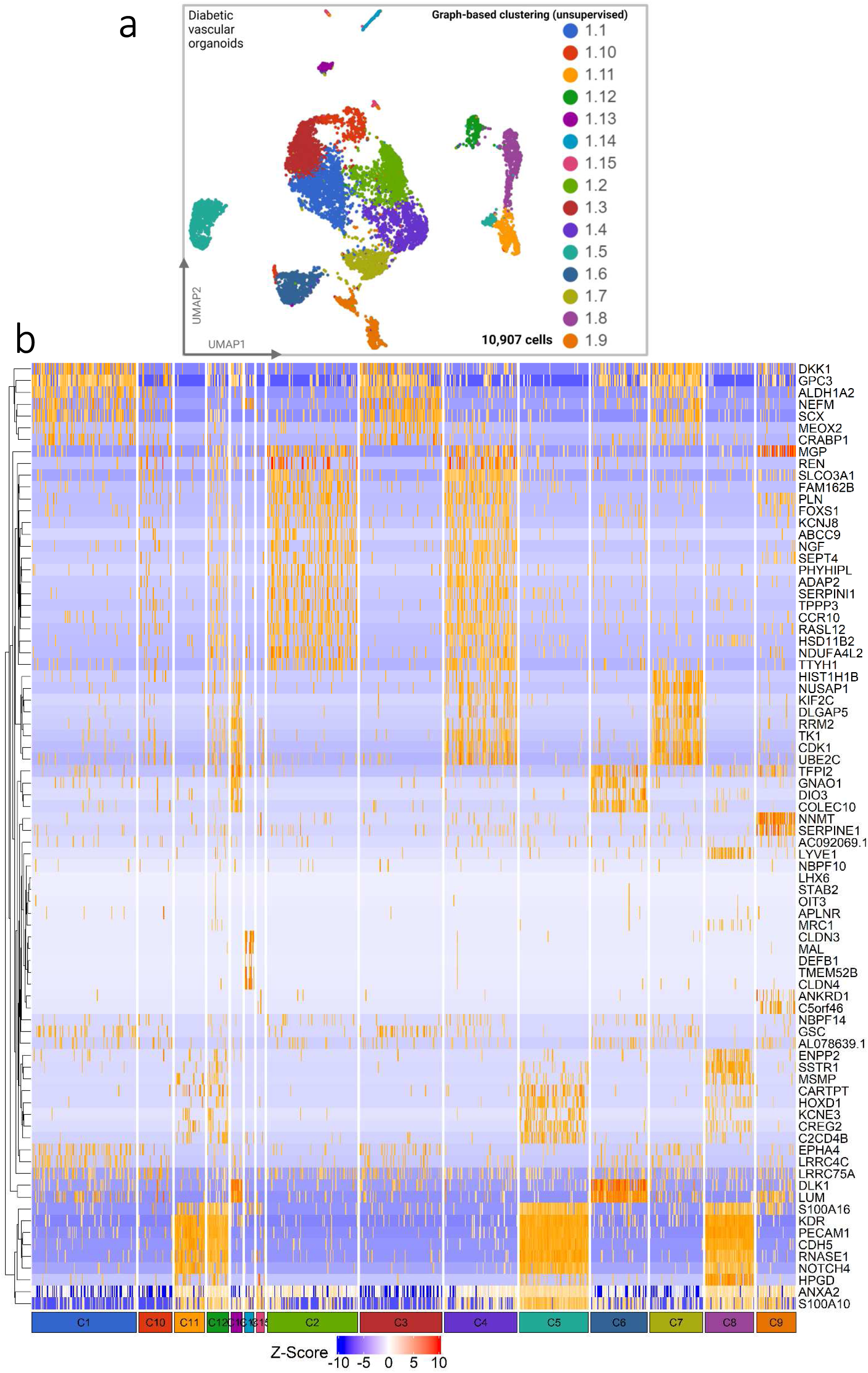
Diabetic vascular organoids represent 15 clusters. **a**, Unsupervised clustering and UMAP projection of DB-VOs. **b**, Heatmap of top cluster-specific biomarkers for 15 clusters of cells of DB-VOs.

**Extended Data Fig. 10.**
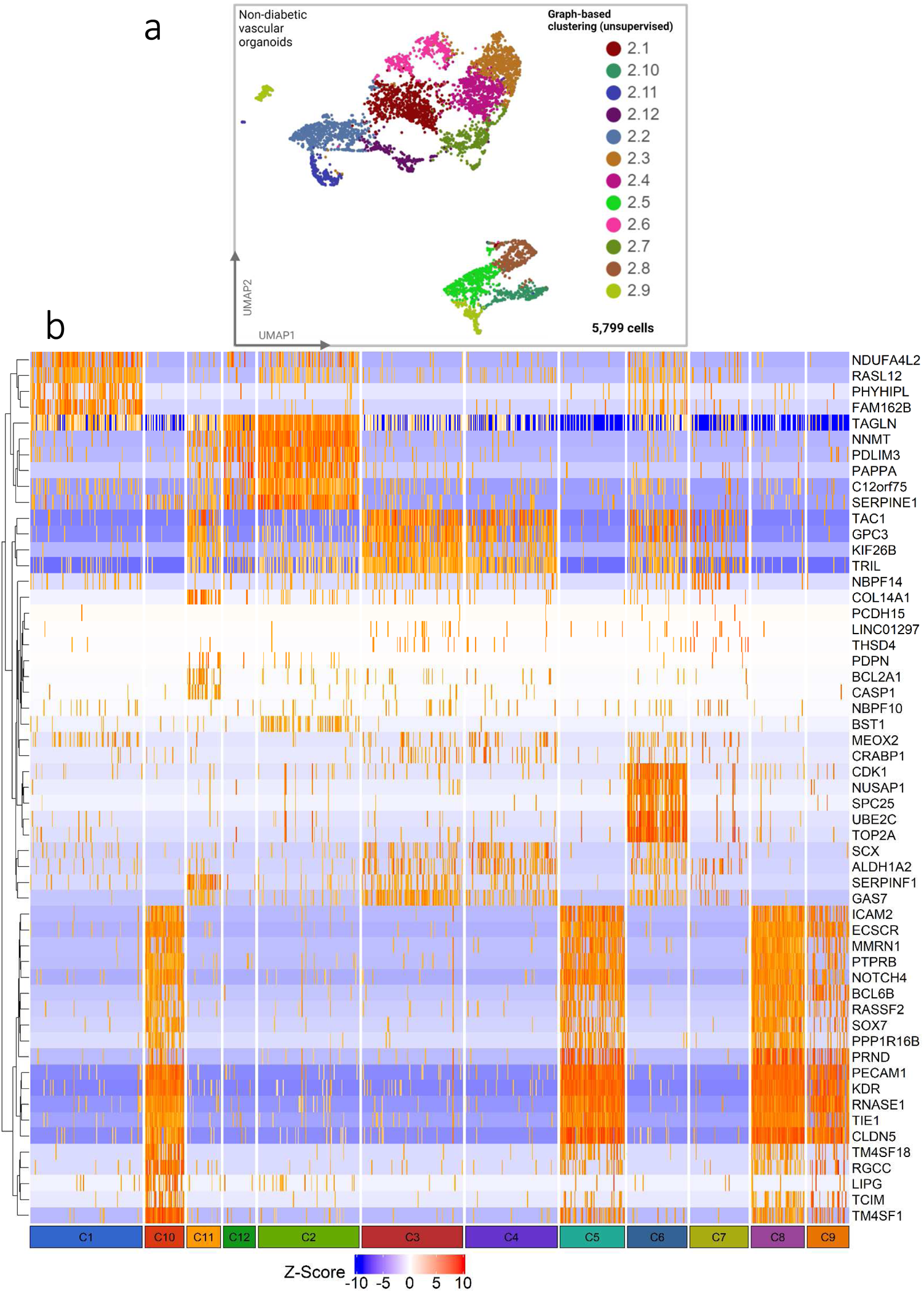
Non-diabetic vascular organoids represent 12 clusters. **a**, Unsupervised clustering and UMAP projection of ND-VOs. **b**, Heatmap of top cluster-specific biomarkers for 12 clusters of cells of ND-VOs.

**Extended Data Fig. 11.**
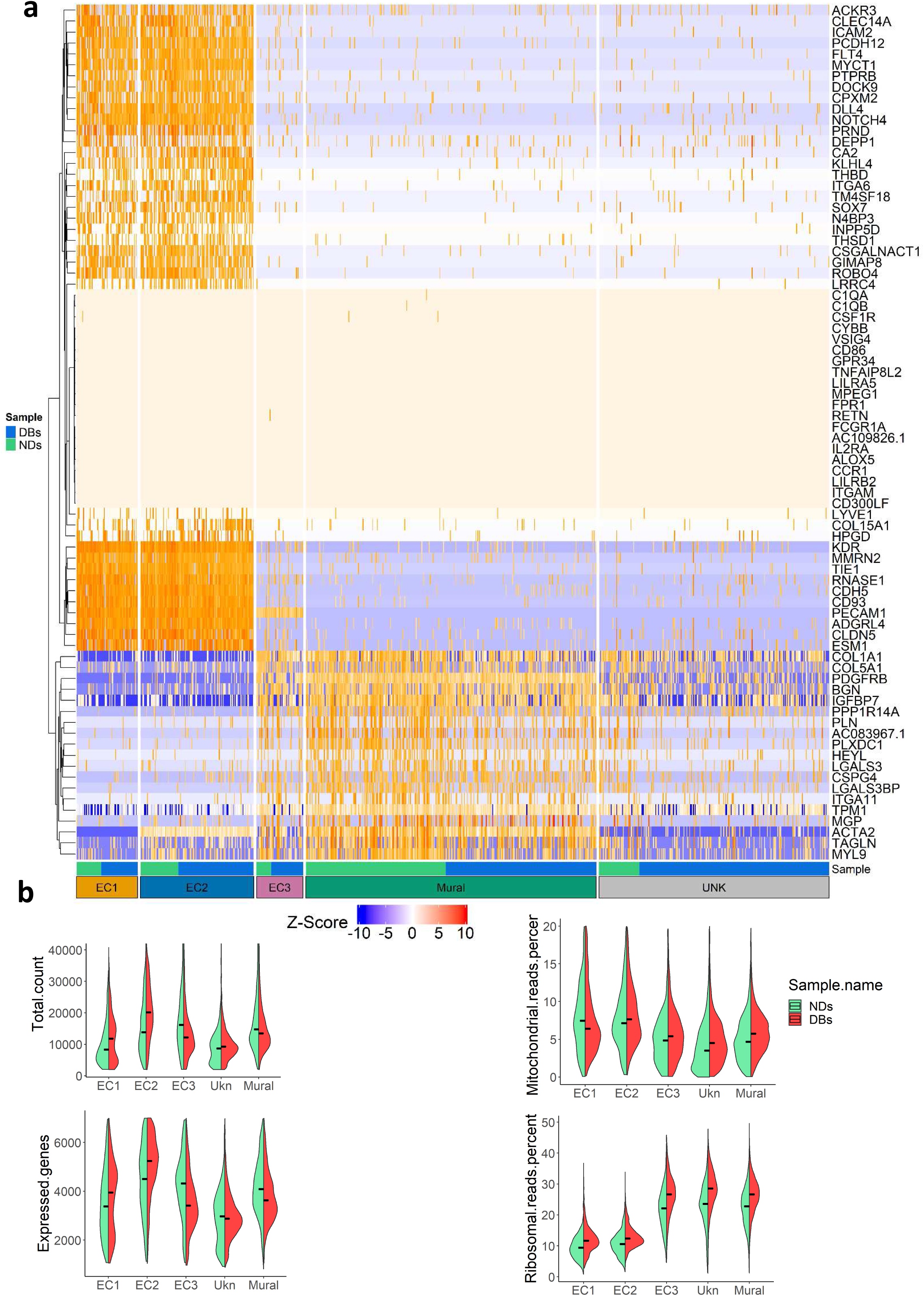
Top cell-specific biomarkers of the subpopulation within vascular organoids. **a**, Heatmap of top cell-specific biomarkers of pooled cells from diabetic and non-diabetic vascular organoids. **b**, Splitviolin showing the number of total counts and expressed genes, and the percent ribosomal reads and mitochondrial reads in each cell type and subpopulation for DB-VOs and ND-VOs.

**Extended Data Fig. 12.**
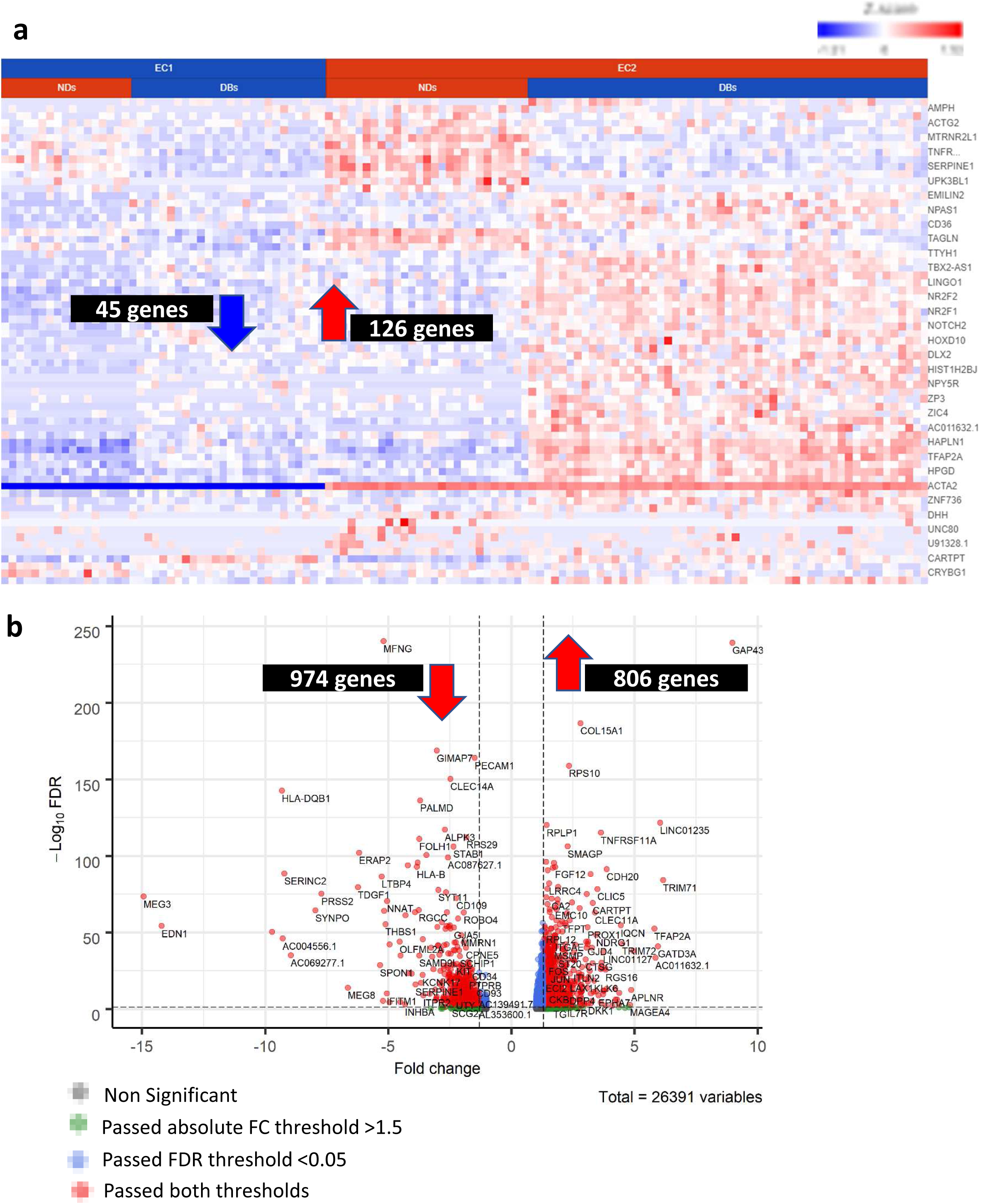
Differentially expressed genes (DEGs) of EC2 versus EC1, and EC2-DB versus EC2-ND. **a**, Heatmap showing top DEGs in EC2 versus EC1 subpopulation of the vascular organoids. In total, 126 genes significantly upregulated in EC2 (FDR < 0.05 & FC > 1.5) and 45 genes significantly downregulated (FDR < 0.05 & FC < -1.5) in EC2. **b**, Volcano plot showing DEGs in EC2-DBs versus EC2-NDs vascular organoids. In total, 806 genes significantly upregulated in EC2-DBs (FDR < 0.05 & FC > 1.5) and 974 genes significantly downregulated (FDR < 0.05 & FC < -1.5) in EC2-DBs. The full list of DEGs were subjected to downstream functional analysis.

## List of supplementary files

**Supplementary Table 1:**
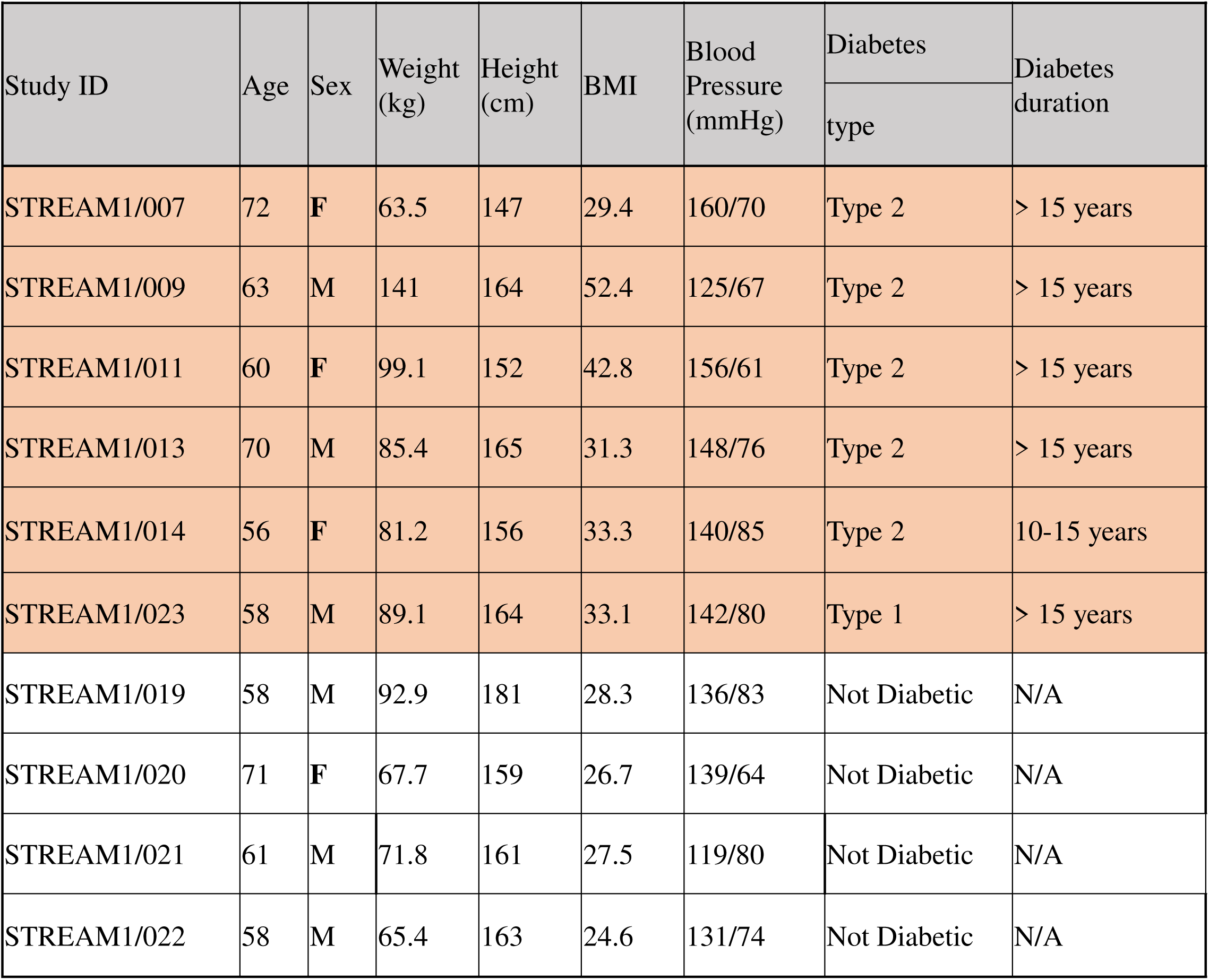
Patient information from which iPS cells were derived, and used in this study.

**Supplementary Video 1:** How to easily cut vascular networks and transfer them back to the ultra-low attachment plate, to reach fully mature organoids within 20 days.

**Supplementary Video 2:** The presence of both small capillaries network and big arteries in the same series of captured confocal images.

**Supplementary Excel sheet 1:** Biomarkers of each cluster by graph-based unsupervised clustering from pooled cells of DB-VOs and ND-VOs (total of 17 clusters).

**Supplementary Excel sheet 2:** Differentially expressed genes of 17 clusters.

**Supplementary Excel sheet 3:** Biomarkers of each cluster by graph-based unsupervised clustering for only of DB-VOs cells (total of 15 clusters).

**Supplementary Excel sheet 4:** Biomarkers of each cluster by graph-based unsupervised clustering for only of ND-VOs cells (total of 12 clusters).

**Supplementary Excel sheet 5:** Biomarkers of each vascular cell type on 2D UMAP.

**Supplementary Excel sheet 6:** Biomarkers of each endothelial subpopulation on 3D UMAP.

**Supplementary Excel sheet 7:** Differentially expressed genes of all cell types in DBs versus NDs.

**Supplementary Excel sheet 8:** Biomarkers of EC2-DB and EC2-ND.

**Supplementary Excel sheet 9:** Merge of unique biomarkers of EC2-DB with differentially expressed genes in EC2-DB versus EC2-ND.

